# Learning from mistakes: Accurate prediction of cell type-specific transcription factor binding

**DOI:** 10.1101/230011

**Authors:** Jens Keilwagen, Stefan Posch, Jan Grau

## Abstract

Computational prediction of cell type-specific, *in-vivo* transcription factor binding sites is still one of the central challenges in regulatory genomics, and a variety of approaches has been proposed for this purpose.

Here, we present our approach that earned a shared first rank in the “ENCODE-DREAM in vivo Transcription Factor Binding Site Prediction Challenge” in 2017. This approach employs features derived from chromatin accessibility, binding motifs, gene expression, genomic sequence and annotation to train classifiers using a supervised, discriminative learning principle. Two further key aspects of this approach are learning classifier parameters in an iterative training procedure that successively adds additional negative examples to the training set, and creating an ensemble prediction by averaging over classifiers obtained for different training cell types.

In post-challenge analyses, we benchmark the influence of different feature sets and find that chromatin accessiblity and binding motifs are sufficient to yield state-of-the-art performance for *in-vivo* binding site predictions. We also show that the iterative training procedure and the ensemble prediction are pivotal for the final prediction performance.

To make predictions of this approach readily accessible, we predict 682 peak lists for a total of 31 transcription factors in 22 primary cell types and tissues, which are available for download at https://www.synapse.org/#!Synapse:syn11526239, and we demonstrate that these may help to yield biological conclusions. Finally, we provide a user-friendly version of our approach as open source software at http://jstacs.de/index.php/Catchitt.

## 1 Introduction

Activation or repression of transcription is one of the fundamental levels of gene regulation. Transcriptional gene regulation depends on transcription factors (TFs), which specifically bind directly to sites in promoters or enhancers of regulated genes or bind indirectly via other, sequence specific TFs. Modeling binding specificities, typically represented as sequence motifs, has been an important topic of bioinformatics since its infancy (Staden, 1984; Berg and von Hippel, 1987). However, it soon became evident that *in-silico* binding site predictions based on sequence motifs alone are insufficient to explain *in-vivo* binding of TFs and that determinants beyond sequence specificity likely play an important role (Stormo and Fields, 1998; Bulyk, 2003).

The emergence of high-throughput techniques like ChIP-chip (Wu *et al*., 2006) or ChIP-seq (Johnson *et al*., 2007) allowed for experimentally determining *in-vivo* TF binding regions on a genome-wide scale. While especially ChIP-seq and derived techniques have the potential to measure TF-specific and cell type-specific binding, the experimental effort and costs currently preclude ChIP-seq experiments for hundreds to thousands of TFs in a variety of different cell types and tissues. Hence, there is a demand for computational methods predicting cell type-specific TF binding with high accuracy. Fortunately, the existence of genome-wide ChIP data for a subset of TFs and cell types also opens up the opportunity to generate more accurate models by supervised machine learning techniques, which may consider further features beyond motif matches.

High-throughput sequencing may also be used to obtain genome-wide assays of chromatin accessibility (e.g., DNase-seq (Hesselberth *et al*., 2009), ATAC-seq (Buenrostro *et al.,* 2013)), which has been considered one of the key features distinguishing functional from non-functional TF binding sites (Galas and Schmitz, 1978; Chen *et al*., 2010). Chromatin accessibility data may yield genome-wide maps of functional binding sites of a large fraction of TFs but, in contrast to ChIP-seq, does not identify the TF binding to a specific region. Hence, the association between bound regions (“footprints”) and TFs is typically derived computationally (Pique-Regi *et al*., 2011).

Following this path, a plenitude of tools (Supplementary Table S1; detailed discussion in Supplementary Text Supplementary Text S1) has been proposed over the last years (e.g., Pique-Regi et al. (2011); Natarajan et al. (2012); Arvey et al. (2012); Luo and Hartemink (2012); Piper et al. (2013); Sherwood et al. (2014); Gusmao et al. (2014); Raj et al. (2015); Kähärä and Lähdesmäki (2015); Kumar and Bucher (2016); Jankowski et al. (2016); Quang and Xie (2017); Liu et al. (2017); Qin and Feng (2017); Schmidt et al. (2017); Chen et al. (2017)). While the general notion of combining sequence signals with chromatin accessibility data and, in some cases, other features is common to the majority of approaches, they differ in several aspects. Specifically, approaches differ in the source of motif information, which may stem from motif databases or from *de-novo* motif discovery. Matches to these motifs are either used as prior information and filtered by their respective DNase-seq signals in a subsequent step, or DNase footprints are first detected and annotated with TFs based on motif matches in those footprints, or, finally, motif and DNase-seq information are processed jointly. Supervised approaches rely on labeled training data, whereas unsupervised approaches may be applied without any a-priorily known binding sites of the TF at hand. Finally, motif and chromatin accessibility data may be complemented with further experimental or computational assays like histone modifications or sequence conservation.

Each of these approaches has its benefits and downsides, and the results of benchmark studies in the respective original publications are ambiguous with regard to their prediction performance. Against this background, the “ENCODE-DREAM in vivo Transcription Factor Binding Site Prediction Challenge” (https://www.synapse.org/#!Synapse:syn6131484) aimed at assessing the performance of tools for predicting cell type-specific TF binding in human using a minimal set of experimental data in a fair and unbiased manner. The challenge setting has advantages over typical benchmark studies, because approaches are typically applied to the challenge data by their authors, ground truth is known only by the challenge organizers, and participants are typically required to provide code and documentation for their method such that predictions can be reproduced.

Participants in the ENCODE-DREAM challenge were allowed to use binding motifs from any source, genomic sequence, gene annotations, *in-silico* DNA shape predictions, and cell type-specific DNase-seq and RNA-seq data. In addition, TF ChIP-seq data and ChIP-seq-derived labels (“bound”, “unbound”, “ambiguous”) were provided for training cell types and training chromosomes. Predictions had then to be made for combinations of TF and cell type not present in the training data on held-out chromosomes. Predictions were evaluated against labels derived from TF ChIP-seq data for that specific TF and test cell type.

Here, we present our approach for predicting cell type-specific TF binding regions earning a shared first rank among 40 international teams, including developers of several established methods (https://www.synapse.org/#!Synapse:syn6131484/wiki/405275). The approach presented in this paper combines several novel ideas. First, we consider motifs from databases, but also motifs learned by de-novo motif discovery from ChIP-seq and DNase-seq data using sparse local inhomogeneous mixture (Slim) models (Keilwagen and Grau, 2015), which may capture short to mid-range intra-motif dependencies. Second, we process DNase-seq data following the binning idea of previous approaches but defining novel statistics computed from the data in those bins, and derive several sequence-based, annotation-based, and RNA-seq-based features. Third, we apply a supervised machine learning approach that employs a discriminative learning principle, which is related to logistic regression but allows for explicit model assumptions with regard to different features. Fourth, discriminative learning is combined with an iterative training approach for refining sets of representative negative examples. Finally, we combine the predictions of classifiers trained in different of these iterations and on different training cell types in an ensemble-like approach.

As this novel approach has already been benchmarked against a large number of competing approaches as part of the ENCODE-DREAM challenge (https://www.synapse.org/#!Synapse:syn6131484/wiki/405275), we focus on the analysis for the contributions of different aspects of this approach on the final prediction performance in this paper. Specifically, we evaluate the contribution of different features, we compare the performance achieved by standard training with that achieved by the iterative training procedure, and we assess the performance of individual classifiers compared with their ensemble prediction. Based on these analyses, we define and benchmark a simplified variant of the proposed approach. Finally, we provide a large collection of predicted, cell type-specific tracks of binding regions for 31 TFs in 22 primary cell types and tissues to make predictions by this approach readily accessible.

## 2 Methods

### 2.1 Data

We use the following types of input data sets as provided by the challenge organizers (https://www.synapse.org/#!Synapse:syn6131484/wiki/402033):

- the raw sequence of the human genome (hg19) and gene annotations according to the gencode v19 annotation (http://www.gencodegenes.org/releases/19.html) (Harrow *et al*., 2012),
- cell type-specific DNase-seq “fold-enrichment coverage” tracks, which represent DNase-seq signal relative to a pseudo control, smoothed in a 150 bp window,
- cell type-specific DNase-seq peak files in “conservative” (IDR threshold of 10% in pseudo replicates) and “relaxed” (no IDR threshold) flavors,
- cell type-specific TPM values from RNA-seq experiments in two bio-replicates for all gencode v19 genes as estimated by RSEM (Li and Dewey, 2011),
- cell type-specific and TF-specific ChIP-seq peak files in “conservative” (IDR threshold of 10% in pseudo replicates) and “relaxed” (no IDR threshold) flavors,
- cell type-specific and TF-specific label files classifying genome-wide 200 bp regions every 50bp into B=“bound”, A=“ambiguous”, and U=“unbound” according to the respective conservative and relaxed ChIP-seq peak files; an overview of the combinations of TF and cell type in the training data, the leaderboard data, and the test data used for evaluation in the final challenge round is given in Supplementary Figure S1.

In addition, we download sequence motifs represented as PWMs from the following collections:

- TF-specific motifs from the databases HOCOMOCO (Kulakovskiy *et al*., 2016) and DBcorrDB (Grau *et al*., 2015a),
- motifs related to epigenetic markers from the epigram pipeline (Whitaker *et al*., 2015).

Details about the motifs considered are given in section "Features" and Supplementary Text S2.

For predicting cell type-specific binding of TFs in additional cell types beyond those considered in the challenge, we download DNase-seq data (FastQ format) from the ENCODE project (encodeproject.org). Specifically, we select all DNase-seq experiments that i) are flagged as “released”, ii) have FastQ files available, iii) are not from immortalized cell lines, iv) have no entry in one of the “Audit error” categories, and v) are not in the “insufficient replicate concordance” category of “Audit not compliant”. A list of the corresponding experiments is obtained from the ENCODE project and experiments are filtered for the existence of at least two replicates, yielding 23 experiments in total. One of these experiments had to be excluded later, because a different DNase protocol with much shorter reads had been used. For the remaining 22 experiments (Supplementary Table S3), all FastQ files are downloaded from ENCODE and processed using ATAC-Seq/DNase-Seq Pipeline (https://github.com/kundajelab/atac_dnase_pipelines, latest git commit: c1d07d38a02af2f0319a69707eee047ab6112ecc (Tue Mar 21 20:31:25 2017)). The data sets are analyzed using the following parameters `-species hg19 -type dnase-seq -subsample 50M -se.` For further analyzes, the relaxed (`./out/peak/idr/pseudo_reps/rep1/*.filt.narrowPeak.gz`) and conservative peaks (`./out/peak/macs2/overlap/*pval0.1*.filt.narrowPeak.gz`) as well as the DNase coverage (`./out/signal/macs2/rep1/*.fc.signal.bigwig`) are used.

In addition, we download ChIP-seq peak files (Supplementary Table S4) matching these cell types and one of the TFs considered. Based on the “relaxed” (i.e., “optimal idr thresholded peaks”) and “conservative” (i.e., “conservative idr thresholded peaks”) peak files, we derive labels for 200 bp windows every 50 bp as proposed for the challenge. Specifically, we label each 200 bp region overlapping a conservative peak by at least 100bp as “bound”. Of the remaining regions, all regions that overlap a relaxed peak by at least 1 bp are labeled “ambiguous”, while all other regions are labeled “unbound”. For a subset of TFs, no conservative peaks are available due to the lack of replicates. In such cases, we also use the relaxed peaks to assign “bound” labels.

### 2.2 Binning the genome

As the final prediction is requested for overlapping 200 bp regions with an offset of 50 bp, we decide to compute features with a matching resolution of 50 bp. To this end, the genome is divided into non-overlapping bins of 50 bp. Features are then either computed directly with that resolution (where possible, e.g., distance to the closest TSS), or first computed with base-pair resolution and afterwards summarized as aggregate values (minimum, maximum, median, or similar statistics) for each 50 bp bin. An odd number of several, adjacent bins, i.e., the respective feature values (see below), is then considered as input of the classifier composed of statistical models for the training process as well as for making predictions. Conceptually, the classifier uses the information from those bins to compute a-posteriori probabilities *P_i_* that center bin *i* (i.e., the central bin of those adjacent bins considered, cf. Figure 1) contains a peak summit. The number of adjacent bins considered is determined from the median across cell types of the median peak widths of a given TF in the individual training cell types.

**Figure 1:**
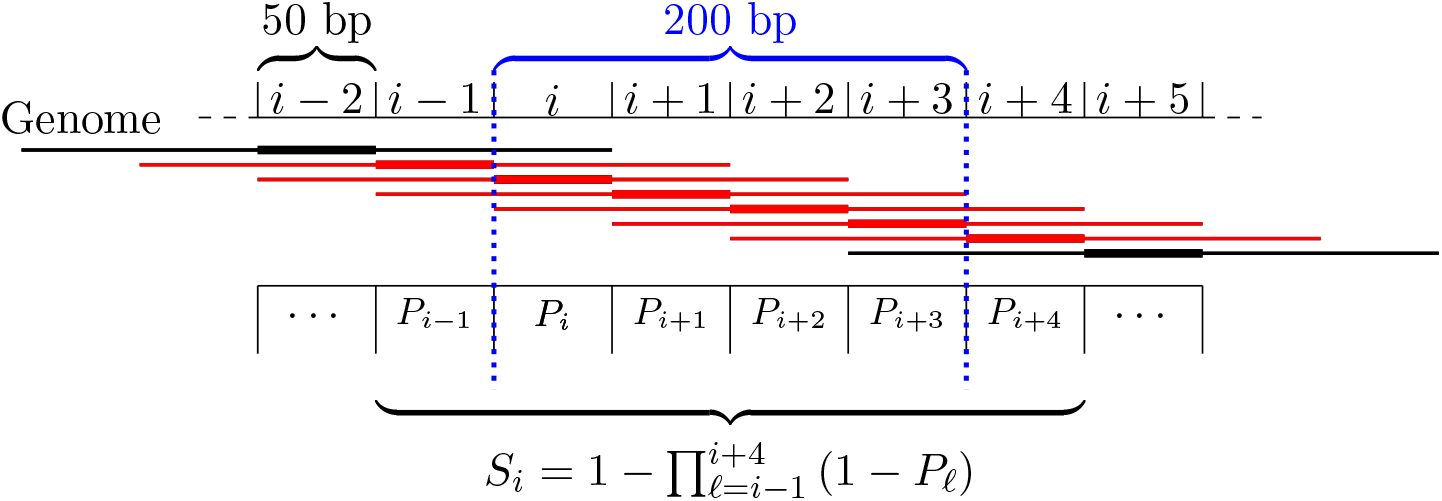
Schema for computing probabilities for 200 bp regions overlapping with predicted peaks spanning five bins in this example. Center bins are indicated by thick lines. Putative peaks are annotated with the probability *P_i_* of being a true peak. All peaks marked in red overlap the region of interest (dotted blue lines) by at least 100 bp and are considered for the prediction. The prediction *S_i_* for the 200 bp region is then computed as the probability that this region overlaps with at least one of the peaks.

### 2.3 Features

The set of features considered may be roughly classified by the source of information: DNase-seq data, motif profiles, raw sequence, RNA-seq data. Here, we give a brief overview of these features, while we provide a complete list of definitions of all features in Supplementary Text S2.

The most informative features with regard to the challenge task are likely motif-based and chromatin accessibility-based features. For obtaining a broad set of binding motifs for each TF at hand, we combine motifs from databases with motifs obtained by de-novo motif discovery from the challenge data. We retrieve PWM models of the TF at hand from the databases HOCOMOCO (Kulakovskiy et al., 2016) and DBcorrDB (Grau *et al*., 2015a). We perform de-novo motif discovery with Dimont (Grau *et al*., 2013) learning PWM and LSlim(3) models (Keilwagen and Grau, 2015) on the “conservative” and “relaxed” ChIP-seq peak files, and also based on the peak files obtained from DNase-seq experiments. In addition, we obtain motifs from the epigram pipeline (Whitaker *et al*., 2015), which are related to DNA methylation and histone marks of active promoters and enhancers. For a specific combination of cell type and TF, we also consider motifs of a set of “peer” motifs, which are determined from the literature (Factorbook, Wang *et al*. (2012)) and by comparing the overlaps between the respective peak lists.

All of these motifs are then used in a sliding window approach to obtain base-pair resolution score profiles, which are summarized by aggregate statistics representing the binding affinity to the strongest binding site (i.e., the maximum log-probability in a bin according to the motif model) as well as general affinity to broader regions (i.e, the logarithm of the average probability in a bin). The set of motifs may comprise models of general binding affinity of the TF at hand but may also capture cell type-specific differences in the binding regions, which could be caused by interaction with other TFs including competition for similar binding sites.

DNase-seq-based features are computed from the “fold-enrichment coverage” tracks and DNase-seq peak files provided with the challenge data. These features quantify short and long range chromatin accessibility, stability of the DNase signal in the region of interest and across different cell types, and overlaps with DNase-seq peak regions.

The set of sequence-based features comprises the raw sequence (i.e., in 1 bp resolution) around the center bin and several measures computed from this sequence, for instance G/C-content, the frequency of CG di-nucleotides, or the length of homo-polymer tracts. Based on the gencode v19 genome annotation, we additionally define features based on overlapping annotation elements like CDS, UTRs, or TSS annotations and based on the distance to the closest TSS annotation in either strand orientation. All of these features are neither cell type-specific nor TF-specific. However, they may represent general features of genomic regions bound by TFs (like CpG islands, GC-rich promoters, or preference for non-coding regions), which might be helpful to rule out false positive predictions based on TF-specific features like motif scores. In addition, the model parameters referring to those features may be adapted in a TF-specific and cell type-specific manner, which may yield auxiliary information for cell type-specific prediction of TF binding as well.

Finally, RNA-seq data are represented by the TPM value of the gene closest to the bin of interest as well as measures of stability within biological replicates and across different cell types.

DNase-seq and RNA-seq-based features are cell type-specific but not TF-specific by design. However, model parameters may adapt to situations where one TF preferentially binds to open chromatin, whereas another TF may also bind in nucleosomal regions.

Feature values are computed using a combination of Perl scripts and Java classes implemented using the Java library Jstacs (Grau et al., 2012). Genome wide feature values with bin-level resolution are pre-computed and stored in a sparse, compressed text format.

### 2.4 Model & basic learning principle

We model the joint distribution of these different features by a simple product of independent densities or discrete distributions (Supplementary Text S3). Specifically, we model numeric features (e.g., DNase-based statistics, motif scores, RNA-seq-based features) by Gaussian densities, discrete, annotation-based features by independent binomial distributions, and raw sequence by a homogeneous Markov model of order 3. All distributions are in the exponential family and parameterized using their natural parameterization (Bishop, 2006; Keilwagen et al., 2010), which allows for unconstrained numerical optimization.

As learning principle, we use a weighted variant (Grau, 2010) of the discriminative maximum conditional likelihood principle (Roos et al. (2005), Supplementary Text S3), which is closely related to logistic regression but allows for making explicit assumptions about the distribution of the underlying data.

### 2.5 Prediction schema

In the challenge, final predictions have been requested for 200 bp windows shifted by 50 bp along the genome, while the proposed classifier predicts a-posteriori probabilities that the current center bin contains a peak summit. To yield the predictions requested, we use these original prediction values (cf. section 2.2) to compute the probability that the 200 bp window overlaps at least one predicted peak by at least 100 bp (Figure 1). Assume that we already computed the a-posterior probabilities *P_i_* that center bin *i* contains the summit of a ChIP-seq peak according to the trained model. Further assume that for the current TF, a peak typically spans 5 bins in total, which corresponds to the center bin, and two bins before and two bins after the center bin in our model (cf. regions marked by lines in Figure 1). Putative peaks overlapping the current 200 bp window starting at bin *i* are those with center bins at *i* — 1 to *i* + 4. Hence, the probability *S_i_* that this window overlaps a peak may be computed as the complementary probability of the event that this window overlaps no predicted peaks, which in turn is just the product of the complementary a-posteriori probabilities *P_ℓ_* of these bins.

### 2.6 Initial training data

For training the model parameters by the discriminative maximum condition likelihood principle, we need labeled input data comprising a set of positive (bound) regions and a set of negative (unbound) regions. In general, a training region is represented by a vector of all feature values described in section Features in an odd number of consecutive bins (see section Binning the genome). In case of positive regions, these are centered at the bin containing the peak summit. We include all such regions around the peak summits of the “conservative peaks” for the current TF and cell type as positive regions.

Since we face a highly imbalanced classification problem with rather few ChIP-seq peaks compared with the large number of bins not covered by a peak, and since the inclusion of all such negative regions into the training set would lead to an inacceptable runtime, we decided to derive representative negative regions by three different sampling strategies. All sampling steps are performed stratified by chromosome.

First, we sample on each training chromosome 10 times as many negative regions (spanning an odd number of consecutive bins) as we find positive regions on that chromosome. Center bins are sampled uniformly over all bins not covered by a “relaxed” peak for the same cell type and TF.

Second, we over-sample negative regions with large DNase-seq median values similar to those of positive examples to yield a representative set of negative regions. This is especially important as these will be regions that are hard to classify using DNase-seq based features but are only lowly represented by the uniform sampling schema. The over-sampling is adjusted for by down-weighting the drawn negative examples to the corresponding frequency among all negative regions (see Supplementary Text S4).

Third, we sample negative regions from regions that are ChIP-seq positive for one of the other cell types (if more than one training cell type exists for that TF), but do not overlap a “relaxed peak” in the current cell type. These negative regions are weighted such that the sum of their weights matches the rate of such regions among all putative negative regions. This sampling schema is intended to foster learning cell type-specific properties as opposed to general properties of the binding regions of the current TF. In this case, we sample four times as many negative regions as we have positives.

Together, these three sampling schemas yield an initial set of negative regions, which serve as input of the discriminative maximum conditional likelihood principle in addition to the positive regions. However, in preliminary tests during the leaderboard round of the challenge, we observed that even this non-trivial sampling schema is not fully satisfactory. As testing (a large number of) other sampling schemas seemed futile, we designed an iterative training schema (Figure 2) that is loosely related to boosting (Freund and Schapire, 1996) and successively complements the initial set of negative training regions with further, informative examples.

**Figure 2:**
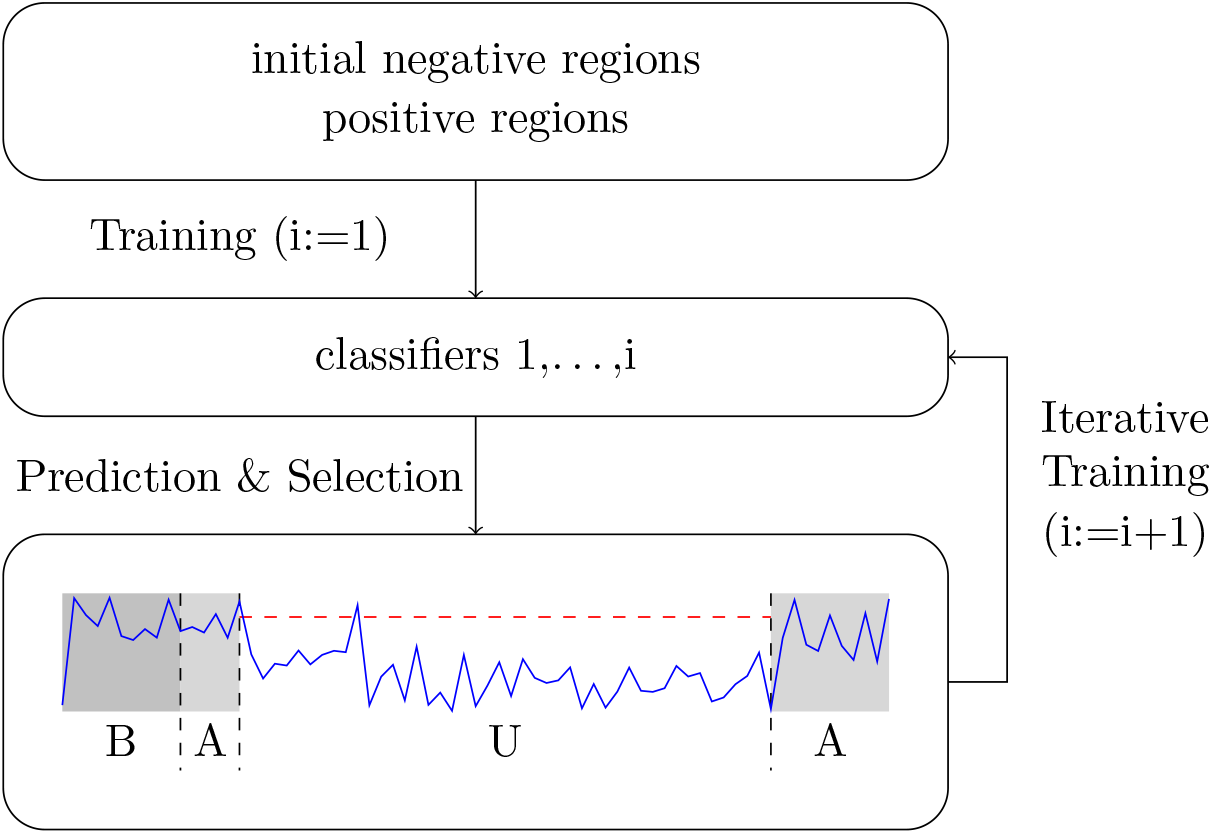
Iterative training procedure. Starting from an initial set of negative regions and the complete set of positive regions, a first classifier is trained, applied to the training data, and putative false positive (i.e, “unbound” regions with large prediction scores) are identified. In each of the subsequent iterations, such regions are added to the set of negative regions, which are in turn used for training refined classifiers. The result of this iterative training procedure is a set of 5 classifiers trained in 5 cycles of the iterative training procedure.

### 2.7 Iterative training

The iterative training procedure is illustrated in Figure 2. Initially, we train a classifier on the negative regions obtained from the sampling schemas explained above and all positive regions. We then use this classifier to obtain a-posteriori probabilities *P_i_* for each bin *i* on training chromosomes. To limit the runtime required for this prediction step, we restrict the prediction to chromosomes chr10 to chr14. These probabilities are then used as input of the prediction schema (section Prediction schema) to yield predictions for the 200 bp regions labeled based on the ChIP-seq training peaks. Given these labels, we may distinguish prediction values of positive regions (label B=“bound”) and negative regions (label U=“unbound”), while regions labeled as A=“ambiguous” are ignored. To select additional negative regions that are likely false positive predictions, we first collect the prediction scores of all positive regions (labeled as B) and determine the corresponding 1% percentile. We then select from the negative regions (labeled as U) all those with a predictions score larger than this 1% percentile, which are subsequently added to the set of negative regions with a weight of 1 per region selected.

In the next iteration, we train a second classifier, again using all positive regions but with negative regions complemented by these additional negative regions. Prediction is then performed using both classifiers, where the predictions of the individual two classifiers (or all previously trained classifiers in subsequent iterations) are averaged per region. Again, regions labeled U with large prediction scores are identified and added to the set of negative regions, which then serve as input of the following iteration. After five rounds of training yielding five classifiers, the iterative training procedure is terminated.

### 2.8 Final prediction

The iterative training procedure is executed for all *K* training cell types with ChIP-seq data for the TF of interest, which yields a total of 5·*K* classifiers. For the final prediction, the prediction schema (section Prediction schema) is applied to all chromosomes and each classifier. These predictions are finally averaged per 200 bp region to yield the final prediction result.

### 2.9 Catchitt: a streamlined open-source implementation

Since the original challenge submission, we have re-implemented the basic approach with the aim of making it more accessible for both, users and developers. Specifically, our objectives were to implement a tool that i) is consolidated into a single runnable JAR file to limit system requirements to a current Java installation only, ii) has an extensible code base eliminating much of the experimental code of the challenge implementation, iii) is applicable to data from individual cell types to reduce data-interdependencies, and iv) may be executed on a standard compute server in acceptable runtime.

To achieve these aims, some parts of the methods have been simplified and streamlined. First, we consider only the most important chromatin accessibility and motif-based features, which reduces runtime and memory consumption. Second, we implement an accelerated motif scanning module that computes whole-genome score profiles even for the complex LSlim models within a few hours. Third, we skip steps that jointly evaluate data and/or feature files from multiple cell types. Specifically, we skip quantile normalization of chromatin accessibility features (although normalization could be performed externally, still), and we omit the sampling step depending on ChIP-seq data for other cell types for determining initial negative regions. We call this implementation “Catchitt” comprising five modules for i) computing chromatin accessiblity features from DNase-seq or ATAC-seq data, ii) computing motif-based features, iii) deriving labels from ChIP-seq peak lists, iv) performing iterative training given feature files and labels, and v) predicting binding probabilities for genomic regions.

### 2.10 Deriving peak lists

For the additional primary cell types and tissues beyond those considered in the challenge, we further process final predictions to yield peak lists in narrowPeak format, which are smaller and easier to handle than the genome-wide probability tracks with 50 bp resolution. To this end, we join contiguous stretches of regions with predicted binding probability above a pre-defined threshold *t* into a common peak region. For each region, we record the maximum probability *p*, and discard bordering regions with a probability below 0.8 · *p*. The resulting regions are then annotated according to the narrowPeak format with a “peak summit” at the center of the region yielding *p*, a “score” of –100 · *log*_10_(1 – *p*), and a “signal value” equal to *p*. We generate “relaxed” peak predictions using *t* = 0.6 and “conservative” peak prediction using *t* = 0.8.

### 2.11 Availability

The original challenge implementation has been developed using the open source Java library Jstacs *(Grau et al.,* 2012) combined with custom Perl and bash scripts for data extraction, conversion, and pipelining. The complete code accompanying the challenge submission is, in accordance with the challenge guidelines, available from https://www.synapse.org/#!Synapse:syn8009967/wiki/412123 including a brief method writeup.

The Catchitt implementation is also based on the Jstacs library and is available as a runnable JAR file at http://jstacs.de/index.php/Catchitt, where we also publish the corresponding source code under GPL 3. Catchitt will be integrated into the Jstacs library with its next release.

## 3 Results

During the ENCODE-DREAM challenge, a large number of approaches created by 40 international teams has been benchmarked on 13 cell type-specific ChIP-seq assays for 12 different TFs in human (Supplementary Figure S1). A set of 109 data sets for the same (and additional) TFs in other cell types was provided for training. Training data comprised cell type-specific DNase-seq data, cell type-specific RNA-seq data, genomic sequence and annotations, and *in-silico* DNA shape predictions. In addition, cell type-specific and TF-specific ChIP-seq data and derived labels were provided for training chromosomes, while predictions were evaluated only on the remaining, held-out chromosomes chr1, chr8, and chr21 that were not provided with any of the ChIP-seq training data. For 200 bp regions shifted by 50 bp, genome-wide predictions of the probability that a specific region overlaps a ChIP-seq peak were requested from the participating teams. Predictions were evaluated by i) the area under the ROC curve (AUC-ROC), ii) the area under the precision-recall curve (AUC-PR), iii) recall at 10% FDR, and iv) recall at 50% FDR on each of the 13 test data sets. These were aggregated per data set based on the average, normalized rank earned for each of these measures in 10 bootstrap samples of the held-out chromosomes, and a final ranking was obtained as the average of these rank statistics (cf. https://www.synapse.org/#!Synapse:syn6131484/wiki/405275).

As a result of this ranking, the approach presented in this paper (team “J-Team”) earned a shared first rank together with the approach created by team “Yuanfang Guan”.

In the following, we investigate the influence of different aspects of the proposed approach on the final prediction performance. First, we inspect the impact of different sets of related features (DNase-seq data, motif scores, RNA-seq data, sequence-based and annotation-based features) on prediction performance. Second, we study the importance of the iterative training approach as opposed to a training on initial training data. Third, we compare the performance of the predictions gained by classifiers trained on training data for individual cell types with the performance of the aggregated prediction obtained by averaging over these cell types. Finally, we apply the proposed method for predicting cell type-specific TF binding for 31 TFs in 22 additional primary cell types yielding a total of 682 prediction tracks.

### 3.1 Impact of feature sets on prediction performance

We use the prediction performance obtained by the proposed approach using all sets of features (section Features), the iterative training procedure (section Iterative training), and the aggregation over all training cell types (section Prediction schema) as a baseline for all further comparisons (Figure 3; “all features”). Throughout this manuscript, we consider AUC-PR as the primary performance measure, since AUC-PR is more informative about classification performance for heavily imbalanced classification problems (Keilwagen *et al*., 2014; Saito and Rehmsmeier, 2015), and recall at the different FDR levels is rather unstable since it corresponds to single points on the precision-recall curve. AUC-PR values are computed using the R-package PRROC (Grau *et al*., 2015b), which has also been used in the ENCODE-DREAM challenge.

**Figure 3:**
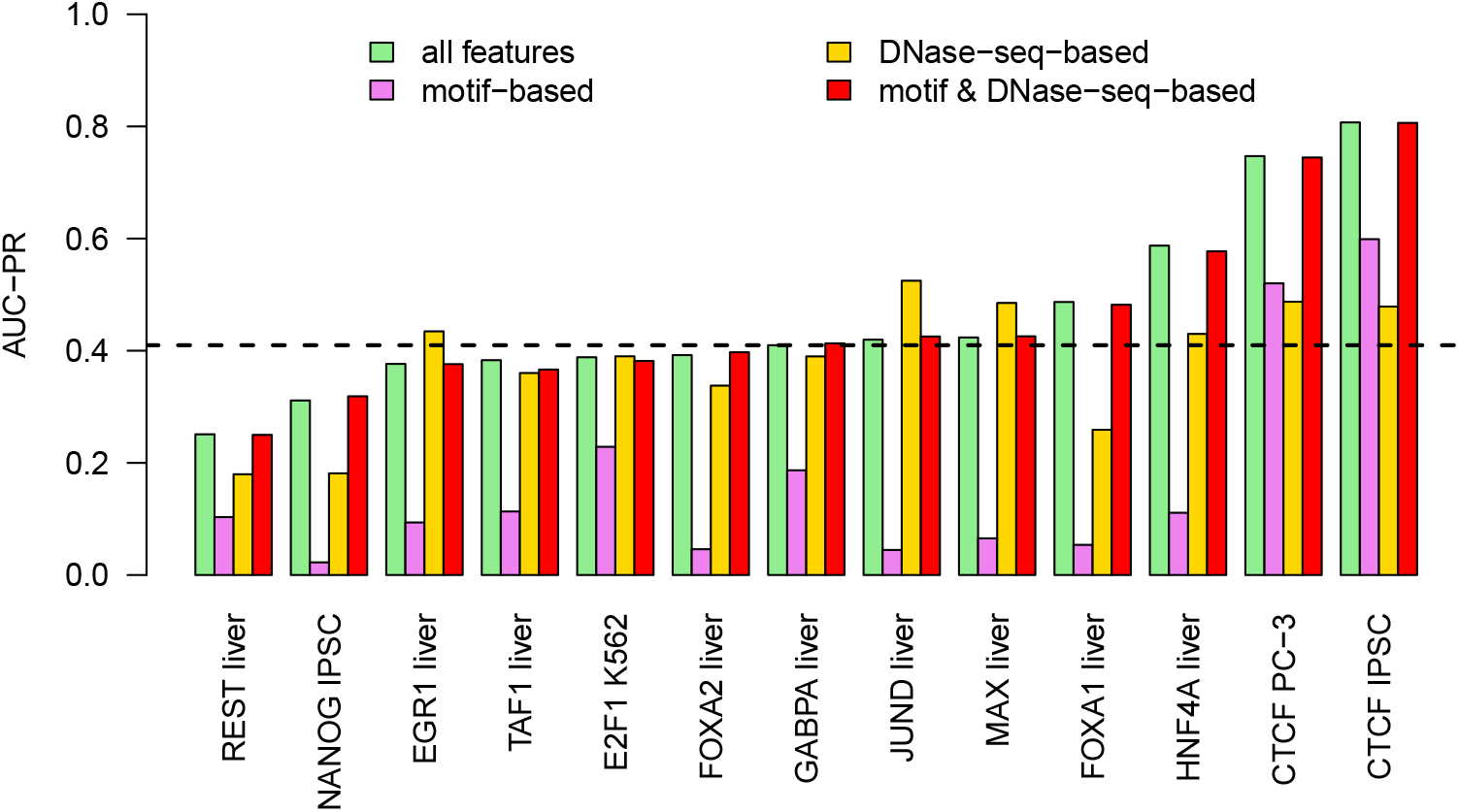
Across cell type performance. For each of the 13 combinations of TF and cell type within the test data, we compute the prediction performance (AUC-PR) on the held-out chromosomes of classifiers i) using all features considered, ii) using only motif-based features, iii) using only DNase-seq-based features, and iv) using only motif-based and DNase-seq-based features. Median performance of classifiers using all features is indicated by a dashed line.

We find that prediction performance as measured by AUC-PR varies greatly among the different transcription factors (Figure 3) with a median AUC-PR value of 0.4098. The best prediction performance is achieved for CTCF, which has a long and information-rich binding motif, in two different cell types (IPSC and PC-3). Above-average performance is also obtained for FOXA1 and HNF4A in liver cells. For most other TFs, we find AUC-PR values around 0.4, whereas we observe a rather low prediction accuracy for NANOG and REST.

To analyze the contribution of selected features on the final prediction performance, we systematically exclude sets of related features from the input data in training and prediction. As a baseline, we measure AUC-PR for the classifier using all feature sets.

In addition, we measure AUC-PR when excluding each individual feature set, where the difference of these two AUC-PR values quantifies the improvement gained by including the feature set (Figure 4).

**Figure 4:**
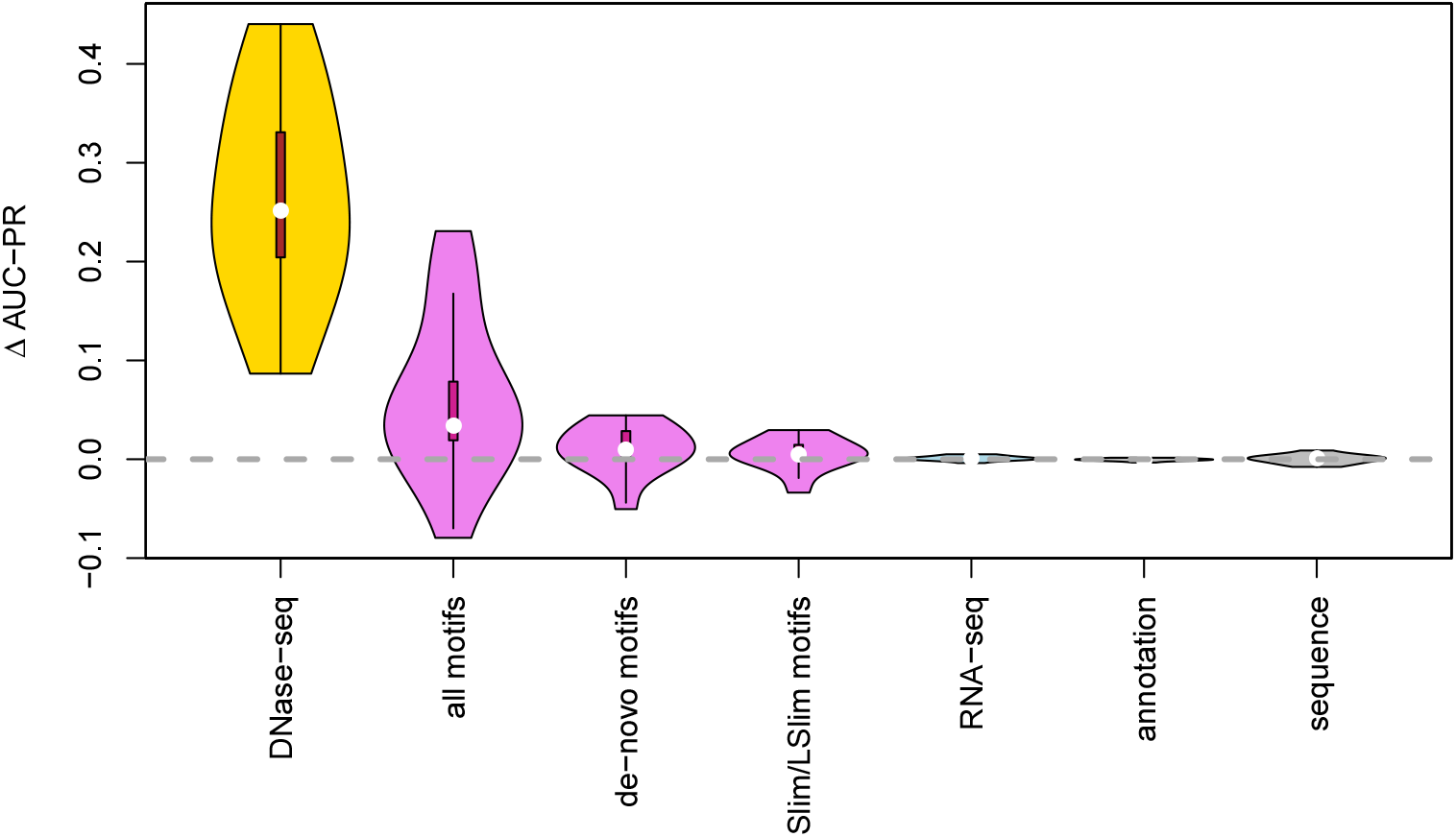
Importance of feature sets. We test the importance of related sets of features by excluding one set of features from the training data, measuring the performance (AUC-PR) of the resulting classifier, and subtracting this AUC-PR value from the corresponding value achieved by the classifier using all features. Hence, if Δ AUC-PR is above zero, the left-out set of features improved the final prediction performance, whereas Δ AUC-PR values below zero indicate a negative effect on prediction performance. We collect the Δ AUC-PR values for all 13 test data sets and visualize these as violin plots.

We observe the greatest impact for the set of features derived from DNase-seq data. The improvement in AUC-PR gained by including DNase-seq data varies between 0.087 for E2F1 and 0.440 for HNF4A with a median of 0.252.

Features based on motif scores (including de-novo discovered motifs and those from databases) also contribute substantially to the final prediction performance. Here, we observe large improvements for some TFs, namely 0.231 for CTCF in IPSC cells, 0.175 for CTCF in PC-3 cells, and 0.167 for FOXA1. By contrast, we observe a decrease in prediction performance in case of JUND (—0.080) when including motif-based features. For the remaining TFs, we find improvements of AUC-PR between 0.008 and 0.079. We further consider two subsets of motifs, namely all motifs obtained by de-novo motif discovery on the challenge data and all Slim/LSlim models capturing intra motif dependencies. For motifs from de-novo motif discovery, we find an improvement for 9 of the 13 data sets and for Slim/LSlim model we find an improvement for 10 of the 13 data sets. However, the absolute improvements (median of 0.011 and 0.006, respectively) are rather small, possibly because i) motifs obtained by de-novo motif discovery might be redundant to those found in databases and ii) intra motif dependencies and heterogeneities captured by Slim/LSlim models (Keilwagen and Grau, 2015) might be partly covered by variations in the motifs from different sources.

Notably, RNA-seq-based features (median 0.001), annotation-based features (0.000), and sequence-based features (0.001) have almost no influence on prediction performance.

Having established that DNase-seq-based and motif-based features have a large impact on prediction performance, we also tested the prediction performance of the proposed approach using *only* features based on DNase-seq data and TF motifs, respectively. We find (Figure 3) that classifiers using exclusively motif-based features already yield a reasonable prediction performance for some TFs (CTCF and, to some extent, E2F1 and GABPA), whereas we observe AUC-PR values below 0.12 for the remaining of TFs. This may be explained by the large number of false positive predictions typically generated by approaches using exclusively motif information, which may only be avoided in case of long, specific motifs as it is the case for CTCF.

Classifiers using only DNase-seq-based features yield a remarkable performance for many of the TFs studied (Figure 3), which is lower than for the motif-based classifier only for the two CTCF datasets. For some datasets (especially JUND but also EGR1, MAX), we even observe that a classifier based on DNase-seq data alone outperforms the classifier utilizing all features.

In case of JUND, the increase in performance when neglecting all non-DNase features can likely be attributed to a strong adaptation of classifier parameters to either cell type-specific binding motifs or cell type-specific co-binding with other TFs, because JUND is the only dataset with an improved performance when excluding motif-based features as discussed above. For all three TFs, we do find an improvement of prediction performance if classifier parameters are trained on the training chromosomes of the test cell type (“within cell type” case; Supplementary Figure S2).

Since DNase-seq-based and motif-based features appear to be the primary feature sets affecting prediction performance, we finally study prediction performance of a classifier using only these two feature sets. We observe that prediction performance using only DNase-seq-based and motif-based features is largely identical to that of the classifier using all features (Figure 3), where we observe the largest loss in AUC-PR for TAF1 (0.017) and the largest gain in AUC-PR for NANOG (0.007). We notice a similar behaviour for the within cell type case (Supplementary Figure S2). As the left-out feature sets include all RNA-seq-based features, this also has the consequence that one cell type-specific assay (namely DNase-seq) is sufficient for predicting TF binding, which broadens the scope of cell types with readily available experimental data that the proposed approach may be applied to.

### 3.2 Iterative training improves prediction performance

As a second key aspect of the proposed approach, we investigate the impact of the iterative training procedure on the final prediction performance. To this end, we compare for each TF the AUC-PR values obtained by averaging over the predictions all five classifiers resulting from the iterative training procedure for all training cell types with the AUC-PR values obtained by only averaging over the initially trained classifiers for all training cell types, i.e., classifiers trained only on the initial training data (section “Initial training data”).

For 11 of the 13 test data sets, we observe an improvement of prediction performance by the iterative training procedure (Figure 5). The largest improvements are achieved for E2F1 (0.114), FOXA2 (0.085), NANOG (0.08), FOXA1 (0.063), and MAX (0.061). Among these are TFs for that we observed a good performance using only DNase-seq-based features (E2F1, MAX) and TFs for which the combination with motif-based features was beneficial (FOXA1, FOXA2, NANOG), which indicates that the additional negative regions added in iterations 2 to 5 do not induce a bias towards either of these two feature types. For four of these five TFs, only one (FOXA2, NANOG, FOXA1) or two (E2F1) training cell types were provided, and the variation between the different classifiers from iterative training may help to avoid overfitting. By contrast, we find a decrease in performance for JUND (0.041) and also TAF1 (0.01), which might be caused by a stronger emphasis on cell type-specific binding regions in subsequent iterations of the iterative training procedure. This hypothesis is also supported by the observation that the iterative training procedure always leads to an increase in prediction performance if classifier parameters are trained on the training chromosomes of the test cell type (Supplementary Figure S3).

**Figure 5:**
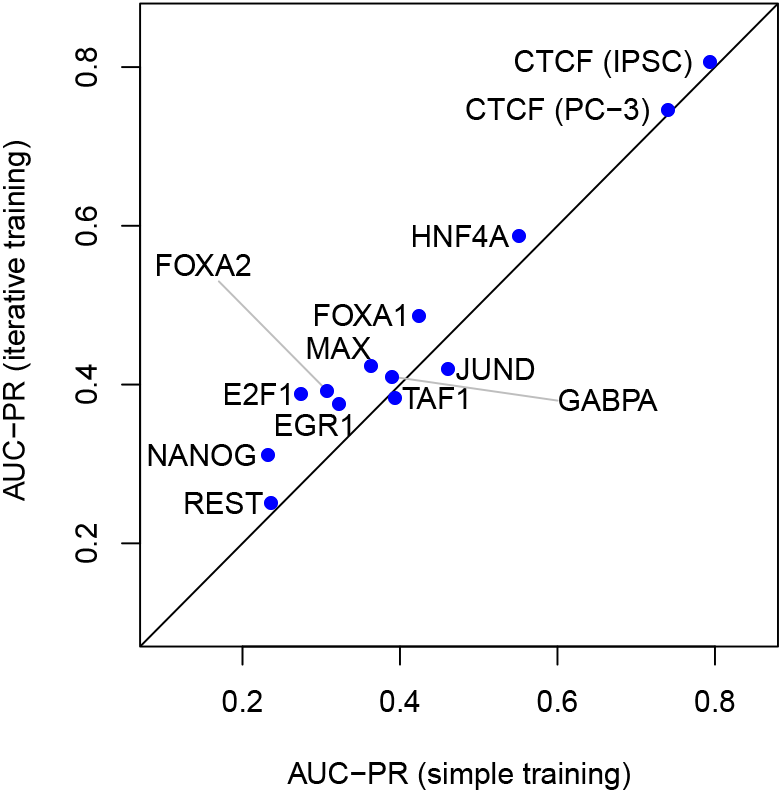
Relevance of the iterative training procedure. For each of the 13 test data sets, we compare the performance (AUC-PR) achieved by the (set of) classifier(s) trained on the initial negative regions (abscissa) with the performance achieved by averaging over all classifiers from the iterative training procedure (ordinate).

### 3.3 Averaging predictions improves over random selection of cell types

For 9 of the 12 TFs considered, data for more than one training cell type is provided with the challenge data. Hence, one central question might be the choice of the cell type used for training and, subsequently, for making predictions for the test cell type. However, the only cell type-specific experimental data available for making that choice are DNase-seq and RNA-seq data, whereas similarity of cell types might depend on the TF considered. Indeed, similarity measures derived from DNase-seq data (e.g., Jaccard coefficients of overlapping DNase-seq peaks, correlation of profiles) or from RNA-seq data (e.g., correlation of TPM values) showed to be non-informative with regard to the similarity of TF binding regions in preliminary studies on the training cell types.

Hence, we consider the choice of the training cell type a latent variable, and average over the predictions generated by the respective classifiers (see section 2.5). As labels of the test cell types have been made available after the challenge, we may now evaluate the impact of this choice on prediction performance and also test the prediction performance of classifiers trained on individual cell types (Figure 6).

**Figure 6:**
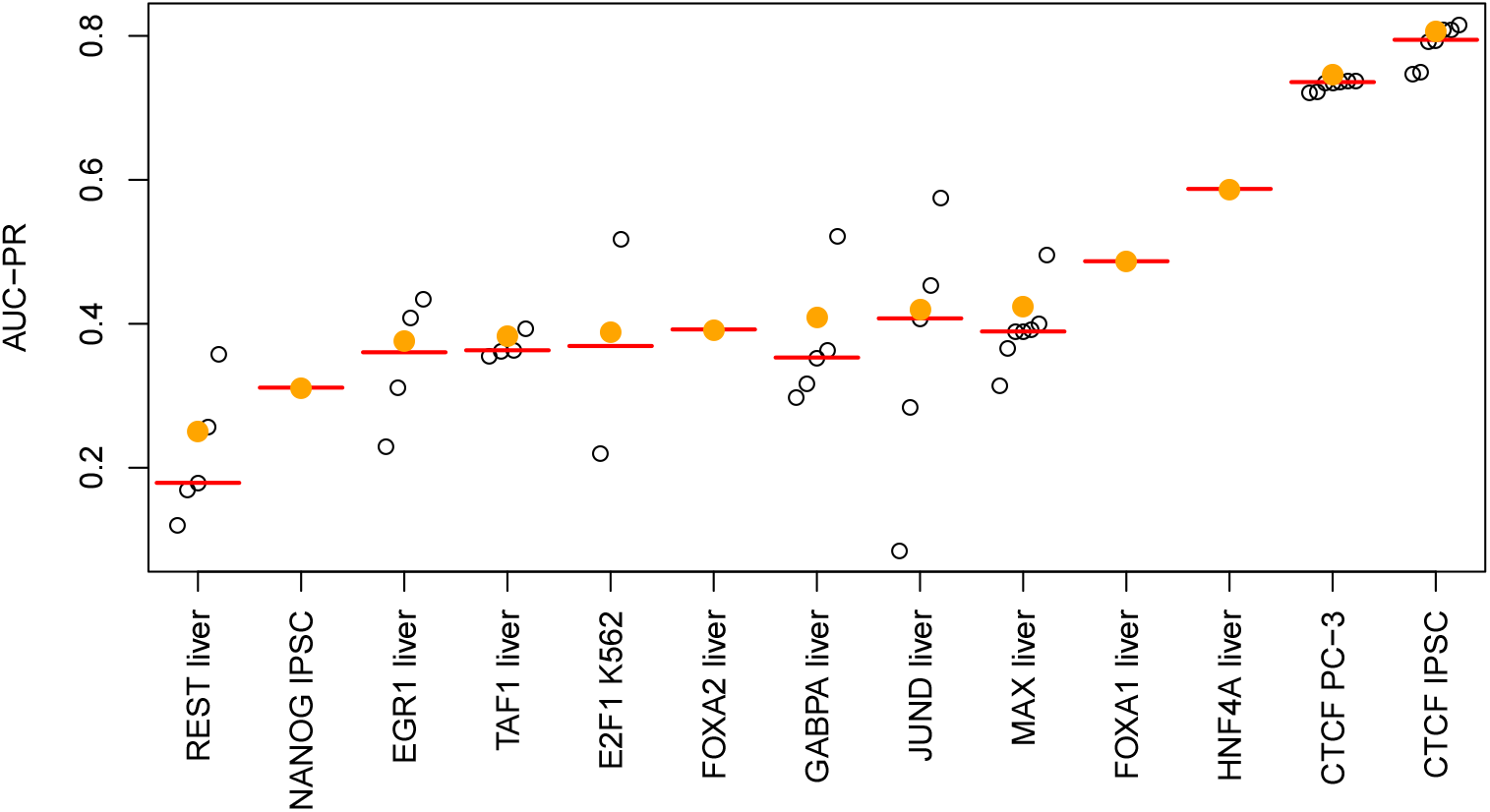
Performance of ensemble classifiers. For each of the 13 test data sets, we compare the performance (AUC-PR) of the individual classifiers trained on single cell types (open circles) to that of the ensemble classifier averaging over all classifiers trained on all training cell types (filled, orange circles). As a reference, we also plot the median of the individual classifiers as a red bar.

For all test data sets with multiple training cell types available, we find that the averaged prediction yields AUC-PR values above the median of the AUC-PR values achieved for individual training cell types. This improvement is especially pronounced for REST, GABPA, and MAX. Hence, we may argue that averaging over the cell type-specific classifiers generally yields more accurate predictions than would be achieved by an uninformed choice of one specific training cell type.

However, we also notice for almost all test data sets with multiple training cell types (the only exception being CTCF for the PC-3 cell type) that the best prediction performance achieved for one of the individual training cell types would have gained, in some cases considerable, improvements over the proposed averaging procedure. Notably, the variance of AUC-PR between the different training cell types is especially pronounced for JUND, which supports the previous hypothesis that some features, for instance binding motifs or co-binding of TFs, are highly cell type-specific for JUND. In general, deriving informative measures of TF-specific cell type similarity based on cell type-specific assays and, for instance, preliminary binding site predictions, would likely lead to a further boost of the performance of computational approaches for predicting cell type-specific TF binding.

### 3.4 Creating a collection of cell type-specific TF binding tracks

Having established that a single type of experimental assay, namely DNase-seq, is sufficient for predicting cell type-specific TF binding with state-of-the-art accuracy, we may now use the classifiers obtained on the training cell types and TFs for predictions on further cell types. To this end, we download DNase-seq data for a collection of primary cell types and tissues (see section Data), process these in the same manner as the original challenge data and, subsequently, extract DNase-seq-dependent features (section Features). We then applied the trained classifiers for all 31 TFs considered in the challenge to these 22 DNase-seq feature sets to yield a total of 682 prediction tracks.

For the selected cell types (Supplementary Table S3), only few cell type and TF-specific ChIP-seq data are available (Supplementary Table S4). On the one hand, this means that the predicted TF binding tracks provide valuable, novel information for the collection of 31 TFs studied. On the other hand, this provides the opportunity to perform benchmarking and sanity checks with regard to the predictions for the subset of these TFs and cell types with corresponding ChIP-seq data available. For benchmarking, we additionally obtain the “relaxed” and (where available) “conservative” peak files from ENCODE and derive the associated labels (“bound”, “unbound”, “ambiguous”) according to the procedure proposed for the ENCODE-DREAM challenge.

For CTCF with ChIP-seq peaks available for multiple cell types, we generally find a prediction performance that is comparable to the performance observed on the challenge data (cf. Supplementary Table S2). For these cell types, AUC-PR values (Supplementary Table S5) range between 0.7720 and 0.8197 if conservative and relaxed peaks are available and if the donors match between the DNase-seq and ChIP-seq experiments, while performance is slightly lower for non-matching donors (0.7322) and in case of missing conservative peaks (0.7270). For JUN, MAX, and MYC, only relaxed peaks are available from ENCODE due to missing replicates. Here, we find AUC-PR values of 0.6310 for JUN, which is substantially larger than for the challenge data, 0.4004 for MAX, which is slightly lower than for the challenge data, and 0.1989 for MYC, which has not been among the test TFs in the challenge but obtained substantially better performance in the leaderboard round.

The 682 genome-wide prediction tracks are still rather large (approx. 880 MB per track) and, hence, demand for substantial storage space that might not be available to the typical user, while the majority of regions are likely not bound by the TF of interest. Hence, we further condense these predictions into predicted peak lists in narrowPeak format by joining contiguous stretches with high binding probability and applying a threshold of 0.6 (relaxed) and 0.8 (conservative) on the maximum probability observed in a predicted “peak”. We provide these peak files for download at https://www.synapse.org/#!Synapse:syn11526239 (doi:10.7303/syn11526239).

To get an impression of the quality of the predicted peaks, we further compute Jaccard coefficients based on peak overlaps (computed using the GenomicRanges R-package (Lawrence *et al*., 2013)) between the predicted peak files and those from the corresponding, available ChIP-seq peaks (Supplementary Tables S6 and S7), and find those to be widely concordant to the previous assessment based on the derived labels.

Based on the predicted peak lists, we may also compare the predicted binding characteristics of the different TFs across cell types. First, we inspect the number of predicted peaks per TF and cell type (Supplementary Figure S4). We find a distinct group of highly abundant TFs (CTCF, GATA3, SPI1, CEBPB, FOXA1, FOXA2, MAX), which typically also show large numbers of peaks in the training data. Among these, we find patterns of cell type specificity from the ubiquitously abundant CTCF to larg-erly varying abundance for GATA3. The remainder of TFs obtains substantially lower numbers of predicted peaks with similar patterns, e.g, for ATF7/ARID3A/NANOG or EP300/TEAD4/JUND, where the latter group has been reported to co-bind in distal enhancers (Xie *et al*., 2013). Next, we study the stability of peak predictions, i.e., the Jaccard coefficients of peaks predicted for each of the TFs in different cell types (Supplementary Figure S5). Again, we find substantial variation among the TFs with GABPA, CTCF, and REST having median Jaccard coefficients above 0.7. Notably, CTCF has been one of the TFs with the largest number of predicted peaks (median 37 455), whereas we observed an order of magnitude less predicted peaks for REST (median 3 364) and GABPA (median 5 430). At the other end of the scale, we find indirectly binding TFs like EP300, or TFs that are highly specific to cell types under-represented in our data like NANOG (stem cells) and HNF4A (liver, kidney, intestines). Finally, we investigate co-binding of TFs by computing the average Jaccard coefficient across cell types for each pair of TFs (Supplementary Figure S6). Here, we observe distinct groups of co-occurring TFs like CTCF/ZNF143 or FOXA1/FOXA2, which are known to interact *in-vivo* (Bailey *et al*., 2015; Ye *et al*., 2016; Motallebipour *et al*., 2009). In addition, we find a larger cluster of TFs with substantial overlaps between their predicted peaks comprising YY1, MAX, CREB1, MYC, E2F6, E2F1, and TAF1. As TAF1 (TATA-Box Binding Protein Associated Factor 1) is associated with transcriptional initiation at the TATA box, one explanation might be that binding sites of these TFs are enriched at core promoters. Indeed, binding to proximal promoters has been reported for MYC/MAX (Guo *et al*., 2014), CREB1 (Zhang *et al*., 2005), YY1 (Li *et al*., 2008), and E2F factors (Rabinovich *et al*., 2008).

### 3.5 Streamlined Catchitt implementation yields competitive performance

We finally compare Catchitt, the simplified implementation of the iterative training approach combining chromatin accessibility and motif scores, to the challenge implementation using DNase-seq-based and motif-based features for the within cell type case. To this end, we select five combinations of cell type and transcription factor spanning the range of performance values observed in the challenge. Specifically, we consider NANOG and TAF1, which obtained the lowest AUC-PR values (cf. Figure S2) for the challenge implementation, CTCF in IPSC cells, which obtained the largest AUC-PR value, and FOXA1 and HNF4A, which obtained medium AUC-PR values but profited substantially from iterative training (cf. Figure S3). We summarize the results of this comparison in Supplementary Table S8. Despite approximately ten-fold reduction in the number of motifs considered and further simplifications (section 2.9), Catchitt still yields competitive AUC-PR values. Ranking the Catchitt results within the original challenge results, we find that performance achieved by Catchitt scores only two ranks lower than the challenge implementation using DNase-seq-based and motif-based features. As before, we find a substantial improvement of prediction performance due to the iterative training procedure.

## 4 Discussion

Predicting *in-vivo* binding sites of a TF of interest *in-silico* is still one of the central challenges in regulatory genomics. A variety of tools and approaches for this purpose have been created over the last years and, among these, the approach presented here is not exceptional in many of its aspects. Specifically, it works on hand-crafted features derived from genomic and experimental data, it considers TF binding motifs and chromatin accessibility as its major sources of information, and it uses supervised learning related to logistic regression. Yet, this approach gained the best performance in the ENCODE-DREAM challenge. Here, we focus on the impact of further, novel aspects of the proposed approach on prediction performance.

With regard to the features considered, we find that motif-based and DNase-seq-based features are pivotal for yielding a reasonable prediction performance for most TFs, while other sequence-based, annotation-based, or RNA-seq-based features have only marginal influence on the prediction result. In case of RNA-seq-based features, however, more sophisticated features than those employed in our approach might have a positive influence on prediction accuracy. In addition, DNA shape might also be informative about true TF binding sites, although *in-silico* shape predictions provided in ENCODE-DREAM are determined based on k-mers, and their influence might also be captured by higher-order Markov models or Slim/LSlim models (Keilwagen and Grau, 2015) employed in the approach presented here.

Previous studies have shown that additional features like sequence conservation (Kumar and Bucher, 2016; Liu *et al*., 2017), histone marks (Pique-Regi *et al*., 2011; Arvey *et al*., 2012; Gusmao *et al*., 2014), or ChIP-seq data of co-factors (Kumar and Bucher, 2016) might also help to predict *in-vivo* TF binding. However, these were not allowed to be used in the ENCODE-DREAM challenge and further experimental assays were unavailable for the training cell types. Hence, we decided to also exclude such features from the studies presented in this paper.

Two aspects of the presented approach, namely the iterative training procedure and aggregation of predictions across training cell types, contribute substantially to the final prediction performance. Both ideas might also be of relevance in related fields. Specifically, the iterative training procedure provides a general schema applicable to imbalanced classification problems, especially when these require sampling of negative examples. In an abstract sense, the aggregation across training cell types corresponds to favoring model averaging over model selection if good selection criteria are hard to find or might yield highly varying results.

Despite its state-of-the-art performance proven in the ENCODE-DREAM challenges, the approach presented here has important limitations. First, the large number of motifs (including those from *de-novo* motif discovery) and DNase-seq-based features lead to high demands with regard to disk space but also runtime, which are likely beyond reach for wet-lab biologists. Disk requirements could be reduced by computing features from (smaller) raw files on demand. However, this would in turn increase running time considerably. Hence, we chose to implement a simplified version of the approach presented here in an open source software available at http://jstacs.de/index.php/Catchitt, which only uses a combination of chromatin accessibility features and motif-based features. In preliminary benchmarks (Supplementary Table S8), this implementation still achieved competitive performance.

Second, the approach proposed here, like any of the other supervised approaches (Natarajan *et al*., 2012; Arvey *et al*., 2012; Luo and Hartemink, 2012; Kahara and Lahdesmaki, 2015; Kumar and Bucher, 2016; Quang and Xie, 2017; Liu *et al*., 2017; Qin and Feng, 2017; Chen *et al*., 2017), requires labeled training data for at least one cell type and the TF of interest to make predictions for this TF in another cell type. While the latter limitation is partly overcome by unsupervised approaches (Pique-Regi *et al*., 2011; Sherwood *et al*., 2014; Gusmao *et al*., 2014; Raj *et al*., 2015; Jankowski *et al*., 2016), this typically comes at the cost of reduced prediction accuracy (Kahara and Lahdesmaki, 2015; Liu *et al*., 2017).

We also provide a large collection of 682 predicted peak files for 31 TFs using 22 DNase-seq data sets for primary cell types and tissues. Benchmarks based on the limited number of available ChIP-seq data indicate that prediction performance on these cell types is comparable to that achieved in the ENCODE-DREAM challenge, where absolute values of AUC-PR measuring prediction accuracy vary greatly between different TFs. For the wide majority of these combinations of TF and cell type, no experimental data about cell type-specific TF binding is available so far, which renders these predictions a valuable resource for questions related to regulatory genomics in these primary cell types and tissues. Preliminary studies raise our confidence that the predicted peak files may indeed help to solve biological questions related to these cell types and TFs.

## Acknowledgements

We would like to express our gratitude to the ENCODE-DREAM organizers, who composed an excellent challenge with clear rules and meaningful performance measures. We would also like to thank Ivan Kulakovskiy, Andrey Lando, and Vsevolod Makeev (team autosome.ru), Wolfgang Kopp (team BlueWhale), Daniel Quang, and Simon van Heeringen for openly sharing their ideas and thoughts during the challenge.

---

**Supplementary Figure S1:**
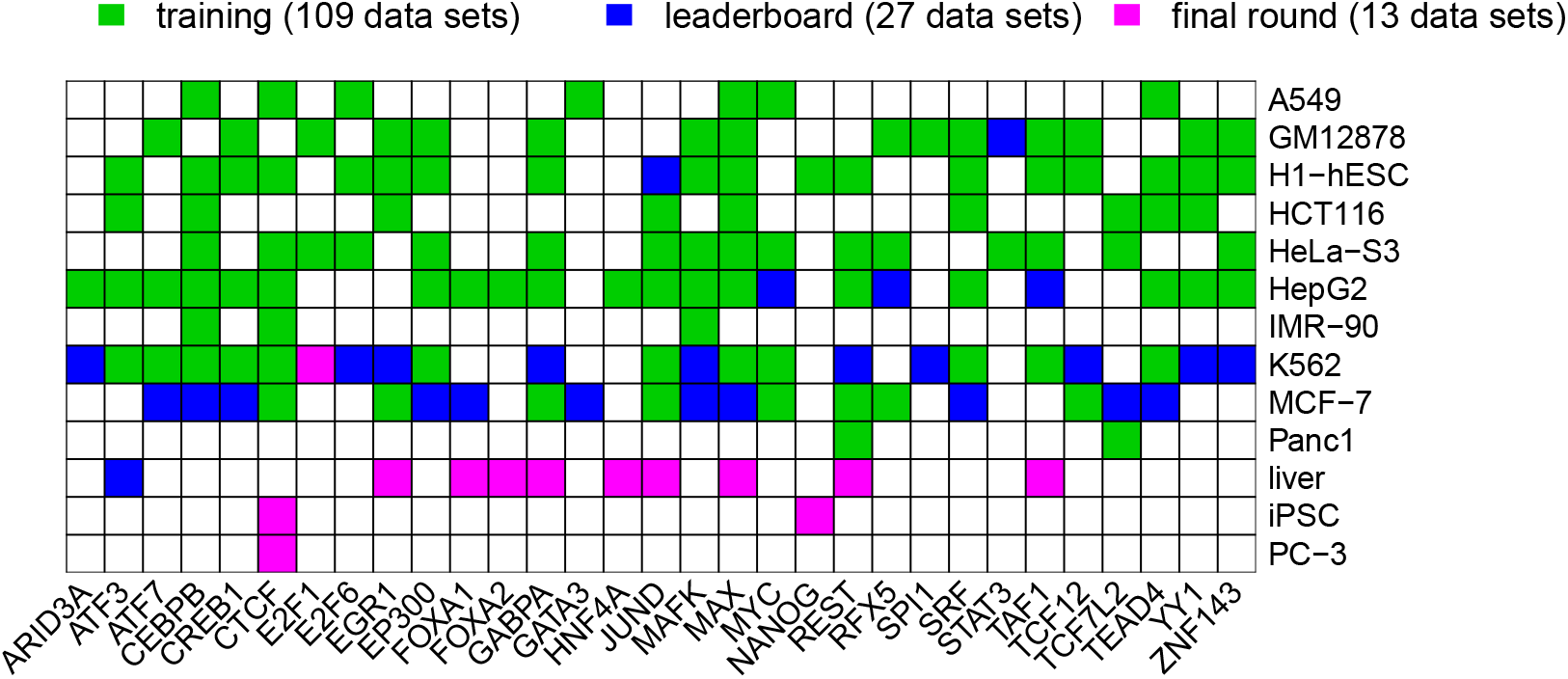
Overview of the combinations of cell type and TF in the ENCODE-DREAM training, leaderboard, and final round sets.

**Supplementary Figure S2:**
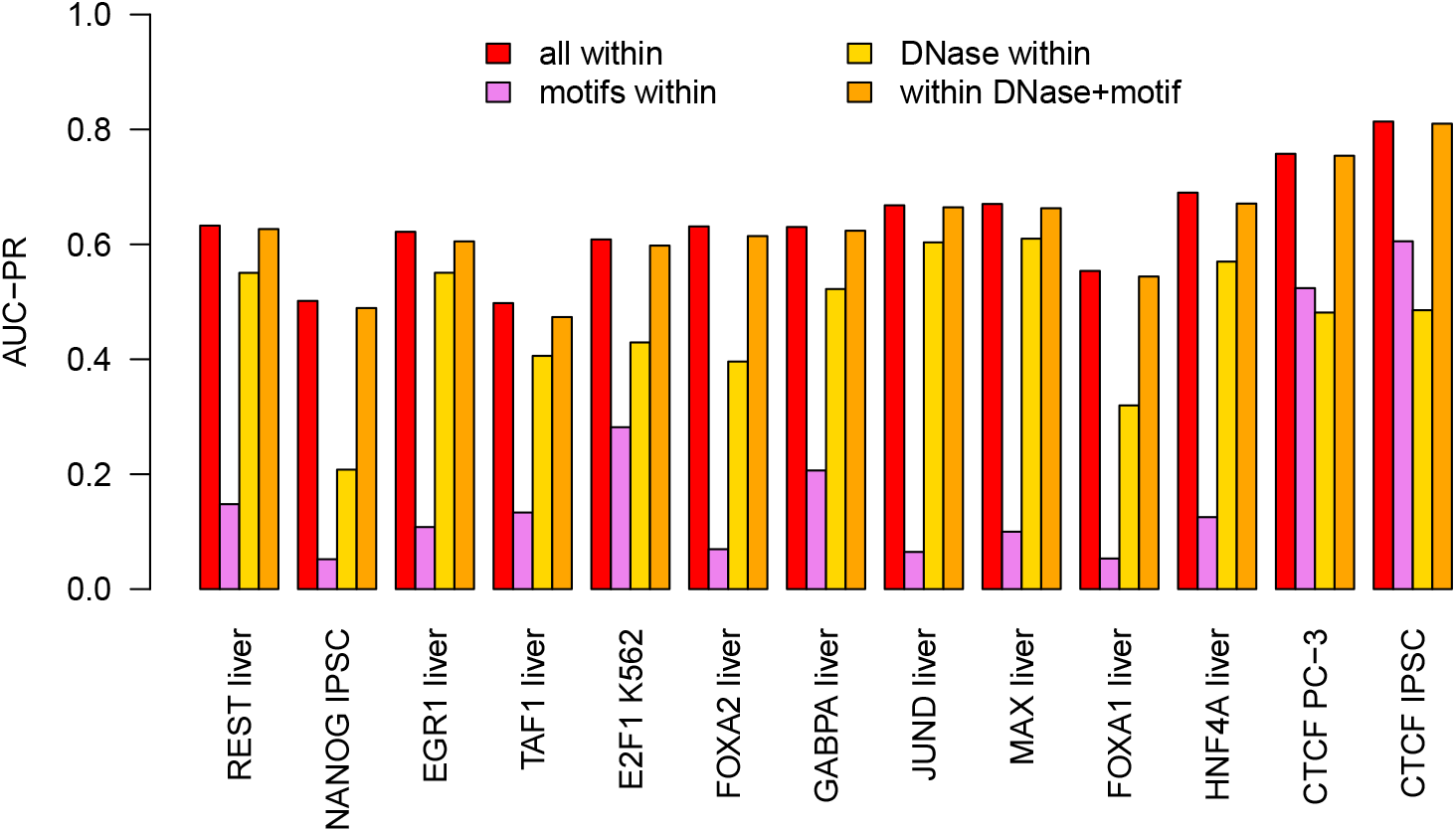
Within cell type performance. For each of the 13 combinations of TF and cell type within the test data, we compute the prediction performance (AUC-PR) on the held-out chromosomes of classifiers i) using all features considered, ii) using only motif-based features, iii) using only DNase-seq-based features, and iv) using only motif-based and DNase-seq-based features. The training data comprises the training chromosomes of the same (test) cell type, while predictions are made for the held-out test chromosomes of that cell type.

**Supplementary Figure S3:**
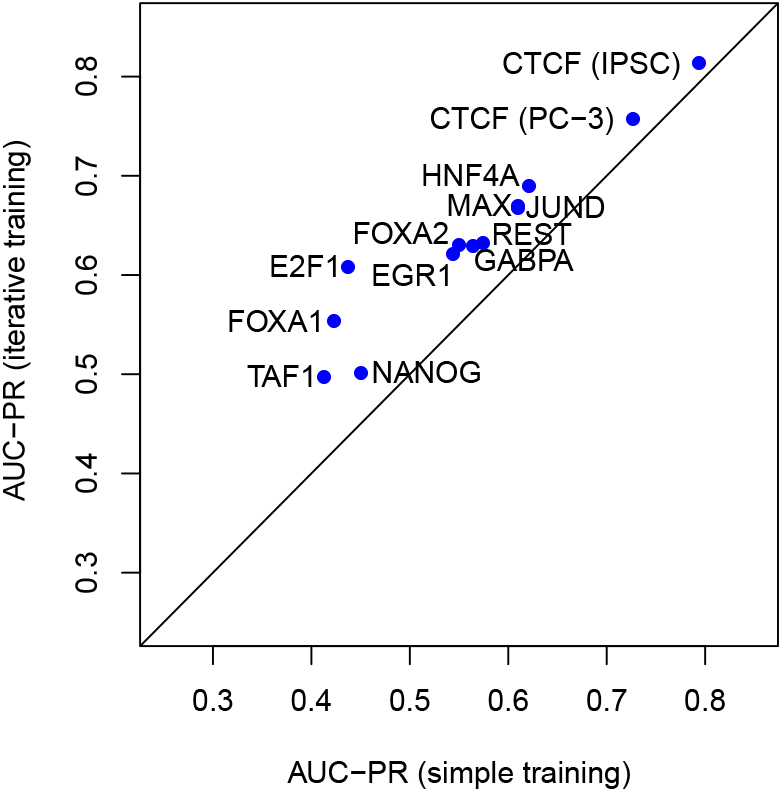
Relevance of the iterative training procedure for within cell type predictions. For each of the 13 test data sets, we compare the performance (AUC-PR) achieved by the (set of) classifier(s) trained on the initial negative regions (abscissa) with the performance achieved by averaging over all classifiers from the iterative training procedure (ordinate). The training data comprises the training chromosomes of the same (test) cell type, while predictions are made for the held-out test chromosomes of that cell type.

**Supplementary Figure S4:**
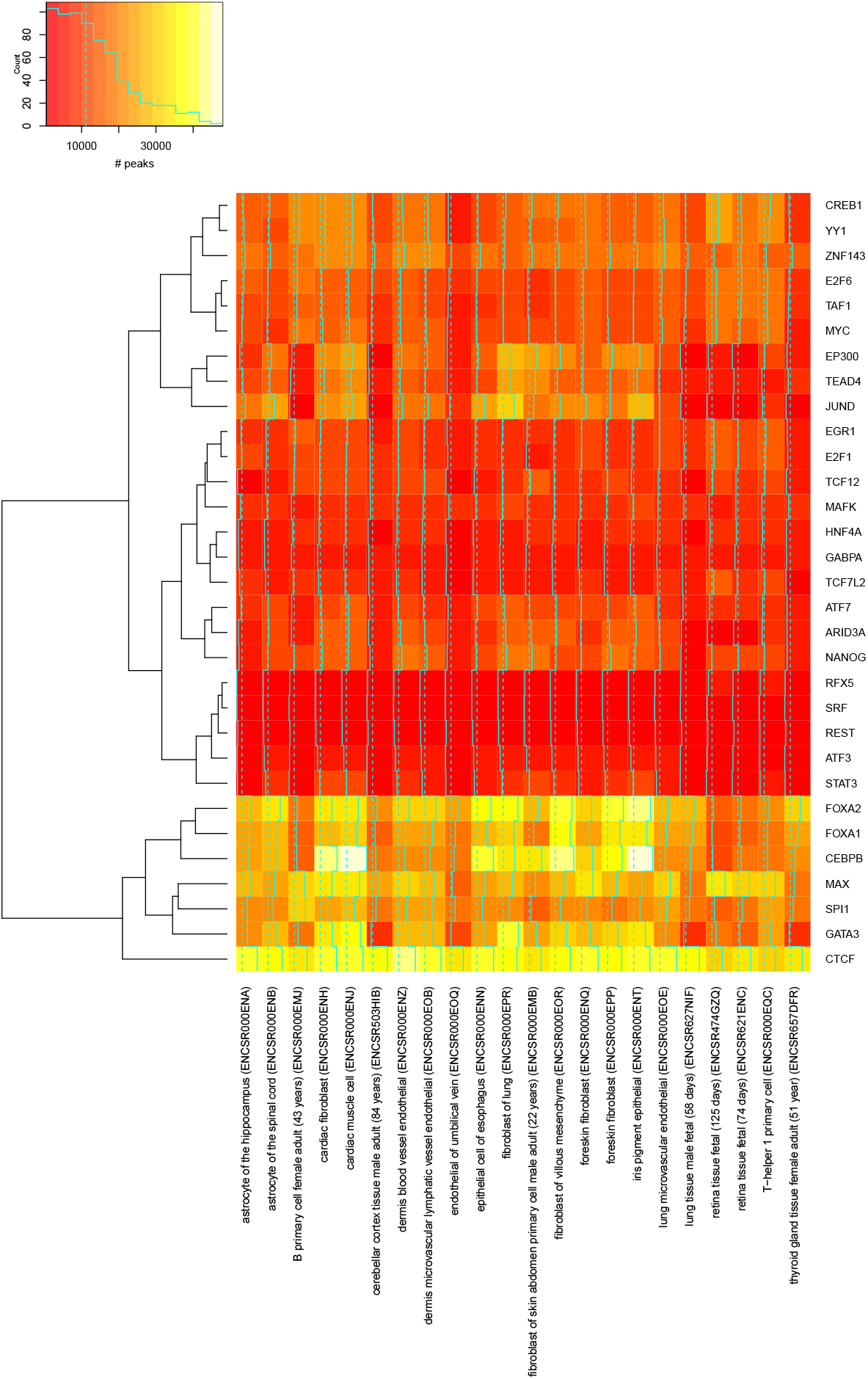
Number of predicted peaks in “conservative” peak files for the studied TFs (rows) in the collection of primary cell types and tissues (columns). In each column of the heatmap, cyan trace lines in addition to colors indicate the corresponding values in each cell. In the color scale, the solid cyan line represents the histogram of values observed in the heatmap. Dashed lines indicate median values across all displayed numbers. Rows are clustered by the `R hclust` function using complete linkage.

**Supplementary Figure S5:**
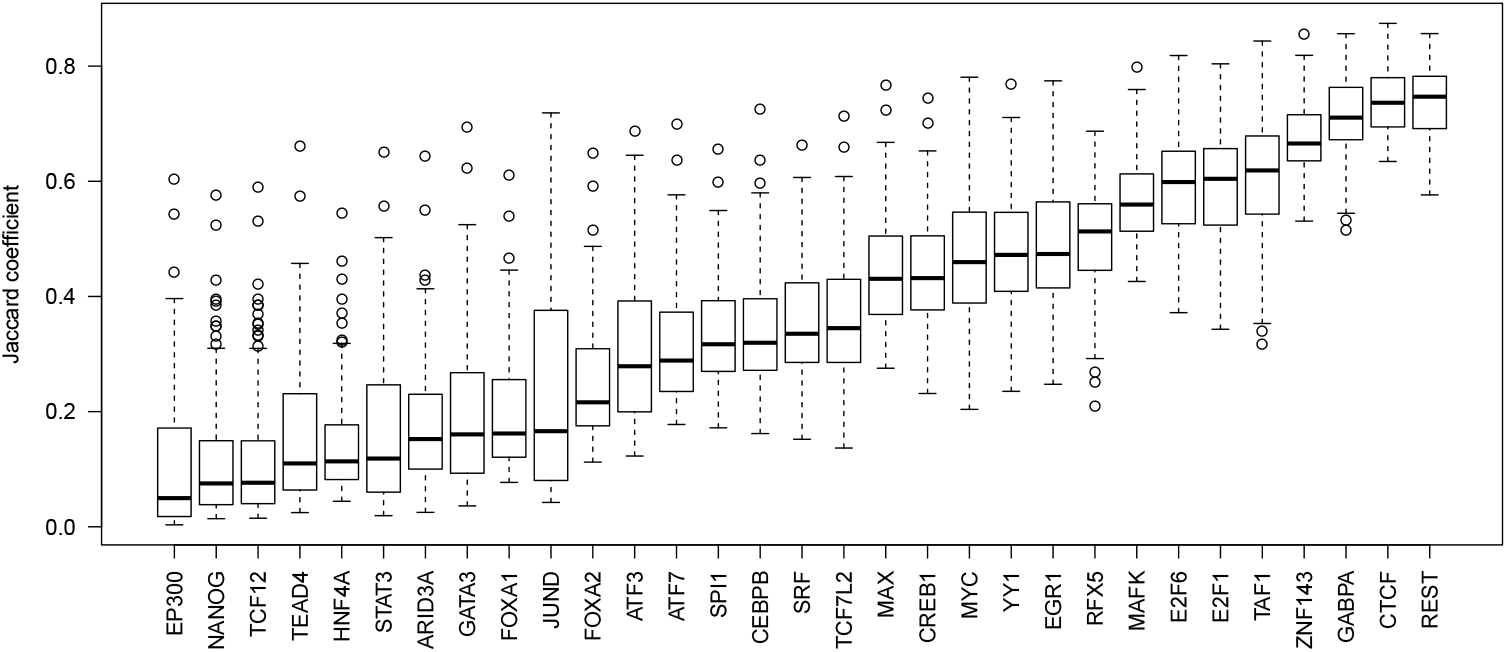
Jaccard coefficients of the different TFs computed on the overlap of the peak files between all pairs of the 22 individual cell types.

**Supplementary Table S1:**
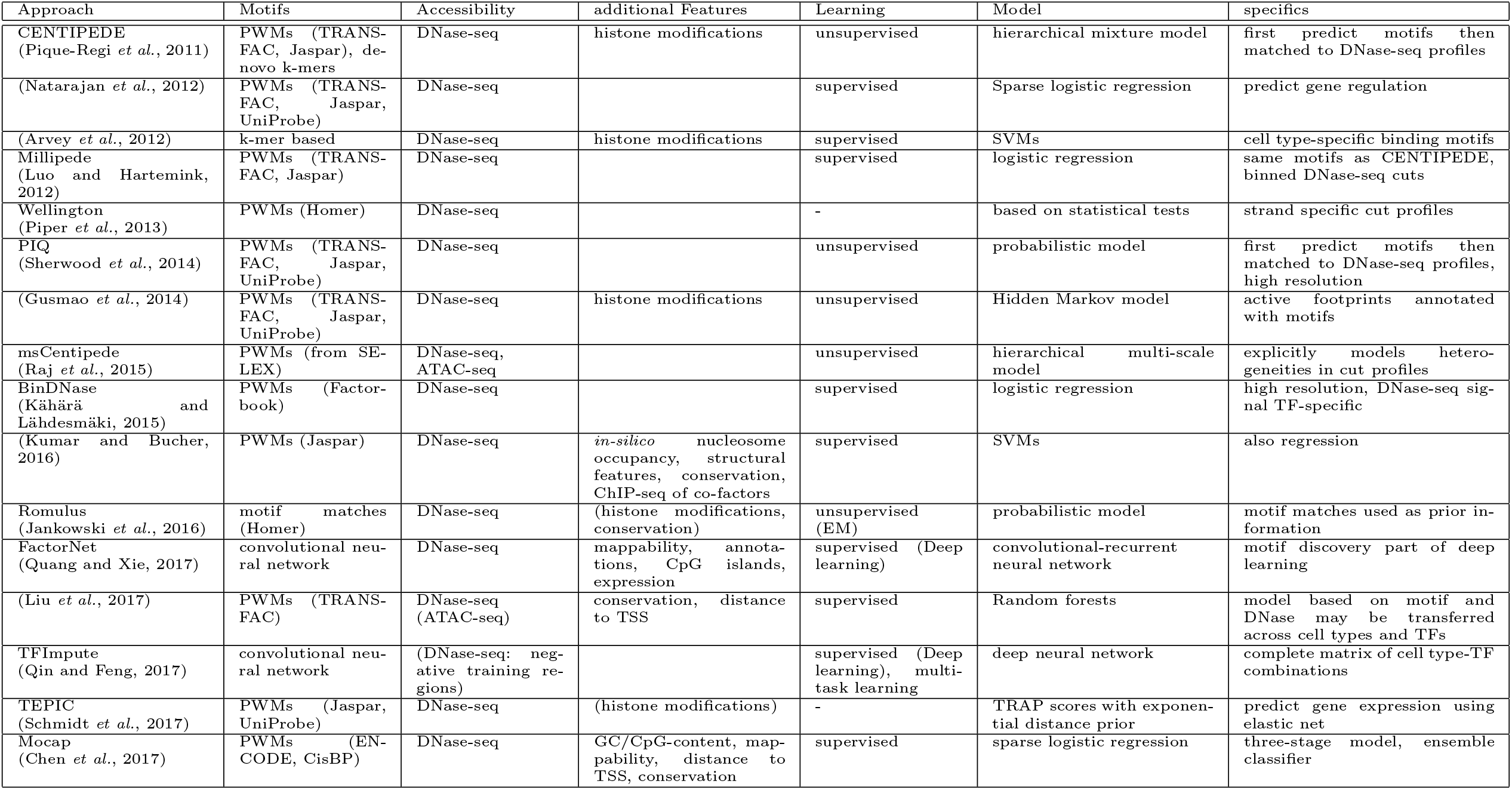
Previous approaches for predicting *in-vivo* transcription factor binding sites and their properties, listed in chronological order.

**Supplementary Table S2:** Performance (AUC-PR) on the test cell types using different sets of features. Columns “all features”, “motif-based”, “DNase-seq-based”, and “motif & DNase-seq-based” correspond to classifiers using only those feature sets, while columns with prefix “w/o” indicate that the given feature set has been excluded when training the classifiers (for details see main text, Figures 3 and 4).

| TF | cell type | all features | motif-based | DNase-seq-based | motif & DNase-seq-based | w/o DNase | w/o motifs | w/o de-novo motifs | w/o Slim/LSlim motifs | w/o RNA-seq | w/o annotation | w/o sequence |
| --- | --- | --- | --- | --- | --- | --- | --- | --- | --- | --- | --- | --- |
| CTCF | IPSC | 0.807 | 0.5989 | 0.479 | 0.806 | 0.6028 | 0.576 | 0.763 | 0.778 | 0.807 | 0.807 | 0.807 |
| CTCF | PC-3 | 0.747 | 0.5202 | 0.487 | 0.745 | 0.5307 | 0.572 | 0.707 | 0.721 | 0.747 | 0.747 | 0.746 |
| E2F1 | K562 | 0.388 | 0.2287 | 0.390 | 0.382 | 0.3017 | 0.326 | 0.366 | 0.382 | 0.384 | 0.390 | 0.385 |
| EGR1 | liver | 0.377 | 0.0937 | 0.435 | 0.376 | 0.1242 | 0.354 | 0.366 | 0.375 | 0.375 | 0.377 | 0.378 |
| FOXA1 | liver | 0.487 | 0.0538 | 0.259 | 0.482 | 0.0713 | 0.321 | 0.458 | 0.478 | 0.487 | 0.488 | 0.482 |
| FOXA2 | liver | 0.392 | 0.0460 | 0.338 | 0.397 | 0.0642 | 0.358 | 0.443 | 0.426 | 0.396 | 0.392 | 0.391 |
| GABPA | liver | 0.410 | 0.1868 | 0.390 | 0.413 | 0.2289 | 0.391 | 0.412 | 0.409 | 0.409 | 0.411 | 0.414 |
| HNF4A | liver | 0.587 | 0.1110 | 0.430 | 0.577 | 0.1471 | 0.509 | 0.573 | 0.573 | 0.586 | 0.586 | 0.579 |
| JUND | liver | 0.420 | 0.0446 | 0.525 | 0.425 | 0.0588 | 0.499 | 0.438 | 0.435 | 0.418 | 0.419 | 0.427 |
| MAX | liver | 0.424 | 0.0654 | 0.485 | 0.426 | 0.0928 | 0.411 | 0.424 | 0.422 | 0.424 | 0.425 | 0.426 |
| NANOG | IPSC | 0.311 | 0.0226 | 0.181 | 0.319 | 0.0304 | 0.291 | 0.306 | 0.306 | 0.313 | 0.312 | 0.317 |
| REST | liver | 0.251 | 0.1033 | 0.180 | 0.250 | 0.1315 | 0.178 | 0.220 | 0.230 | 0.248 | 0.254 | 0.250 |
| TAF1 | liver | 0.383 | 0.1133 | 0.360 | 0.366 | 0.1720 | 0.375 | 0.382 | 0.384 | 0.378 | 0.382 | 0.381 |

**Supplementary Table S3:** Experiment IDs, tissue/cell type information, and biosample “Term ID” of the ENCODE DNase-seq data used in this study. The list of experiments was obtained from https://www.encodeproject.org/report.tsv?type=Experiment&assay_title=DNase-seq&status=released&assembly=hgl9&files.file_type=fastq&audit.NOT_COMPLIANT.category. (accessed March 2, 2017)

| Experiment ID | Donor ID | Tissue/Cell Type | Term ID |
| --- | --- | --- | --- |
| ENCSR000ENA | ENCDO223AAA | astrocyte of the hippocampus | CL:0002604 |
| ENCSR000ENB | ENCDO224AAA | astrocyte of the spinal cord | CL:0002606 |
| ENCSR000ENH | ENCDO095AAA | cardiac fibroblast | CL:0002548 |
| ENCSR000ENJ | ENCDO330AAA | cardiac muscle cell | CL:0000746 |
| ENCSR000ENN | ENCDO104AAA | epithelial cell of esophagus | CL:0002252 |
| ENCSR000ENQ | ENCDO232AAA | foreskin fibroblast | CL:1001608 |
| ENCSR000ENT | ENCDO100AAA | iris pigment epithelial | CL:0002565 |
| ENCSR000EOE | ENCDO238AAA | lung microvascular endothelial | CL:2000016 |
| ENCSR000ENZ | ENCDO241AAA | dermis blood vessel endothelial | CL:2000010 |
| ENCSR000EOB | ENCDO243AAA | dermis microvascular lymphatic vessel endothelial | CL:2000041 |
| ENCSR000EOQ | ENCDO000AAS | endothelial of umbilical vein | CL:0002618 |
| ENCSR000EOR | ENCDO253AAA | fibroblast of villous mesenchyme | CL:0002558 |
| ENCSR000EPP | ENCDO191CQJ | foreskin fibroblast | CL:1001608 |
| ENCSR000EPR | ENCDO269AAA | fibroblast of lung | CL:0002553 |
| ENCSR000EQC | ENCDO334AAA | T-helper 1 primary cell | CL:0000545 |
| ENCSR000EMB | ENCDO442SWC | fibroblast of skin abdomen male adult (22 years) | CL:2000013 |
| ENCSR000EMJ | ENCDO114AAA | B primary cell female adult (43 years) | CL:0000236 |
| ENCSR621ENC | ENCDO539WIJ | retina tissue fetal (74 days) | UBERON:0000966 |
| ENCSR474GZQ | ENCDO225GSN | retina tissue fetal (125 days) | UBERON:0000966 |
| ENCSR503HIB | ENCDO240JUB | cerebellar cortex tissue male adult (84 years) | UBERON:0002129 |
| ENCSR627NIF | ENCDO652XOU | lung tissue male fetal (58 days) | UBERON:0002048 |
| ENCSR657DFR | ENCDO271OUW | thyroid gland tissue female adult (51 year) | UBERON:0002046 |

**Supplementary Table S4:** ChIP-seq data sets available for the primary cell types and tissues. The last seven ChIP-seq data sets provide only “relaxed” peak lists.

| TF | Experiment ID | File ID | Donor ID | Tissue/Cell Type | Type |
| --- | --- | --- | --- | --- | --- |
| CTCF | ENCSR000DSU | ENCFF312HCK | ENCDO224AAA | astrocyte of the spinal cord | relaxed |
| CTCF | ENCSR000DSU | ENCFF787GLH | ENCDO224AAA | astrocyte of the spinal cord | conservative |
| CTCF | ENCSR000DTI | ENCFF266GGD | ENCDO330AAA | cardiac muscle cell | relaxed |
| CTCF | ENCSR000DTI | ENCFF386NQE | ENCDO330AAA | cardiac muscle cell | conservative |
| CTCF | ENCSR000DTR | ENCFF528VFN | ENCDO104AAA | epithelial cell of esophagus | relaxed |
| CTCF | ENCSR000DTR | ENCFF373BXG | ENCDO104AAA | epithelial cell of esophagus | conservative |
| CTCF | ENCSR000DPM | ENCFF681OWQ | ENCDO001AAA | fibroblast of lung | relaxed |
| CTCF | ENCSR000DPM | ENCFF138PXI | ENCDO001AAA | fibroblast of lung | conservative |
| CTCF | ENCSR000DVQ | ENCFF738CXX | ENCDO253AAA | fibroblast of villous mesenchyme | relaxed |
| CTCF | ENCSR000DVQ | ENCFF199ZDU | ENCDO253AAA | fibroblast of villous mesenchyme | conservative |
| CTCF | ENCSR000DWQ | ENCFF337WIE | ENCDO191CQJ | foreskin fibroblast | relaxed |
| CTCF | ENCSR000DWQ | ENCFF275AVH | ENCDO191CQJ | foreskin fibroblast | conservative |
| CTCF | ENCSR000DLW | ENCFF002DBA | ENCDO000AAS | endothelial cell of umbilical vein | relaxed |
| CTCF | ENCSR000DWY | ENCFF002DDO | ENCDO269AAA | fibroblast of lung | relaxed |
| CTCF | ENCSR000DUH | ENCFF002DCY | ENCDO232AAA | foreskin fibroblast | relaxed |
| CTCF | ENCSR000DQI | ENCFF649IRT | ENCDO000AAG | foreskin fibroblast | relaxed |
| JUN | ENCSR000EFA | ENCFF002CVC | ENCDO000AAS | endothelial cell of umbilical vein | relaxed |
| MAX | ENCSR000EEZ | ENCFF002CVE | ENCDO000AAS | endothelial cell of umbilical vein | relaxed |
| MYC | ENCSR000DLU | ENCFF002DAZ | ENCDO000AAS | endothelial cell of umbilical vein | relaxed |

**Supplementary Table S5:** Prediction performance on primary cell types and tissues using labels derived from ChIP-seq data. Here, we include all performance measures considered in the ENCODE-DREAM challenge. *: labels determined from only relaxed peaks.

| TF | DNase ID | Experiment ID | Matching donor | AUC-ROC | AUC-PR | recall @ 10% FDR | recall @ 50% FDR |
| --- | --- | --- | --- | --- | --- | --- | --- |
| CTCF | ENCSR000ENB | ENCSR000DSU | yes | 0.9953 | 0.7895 | 0.5603 | 0.8240 |
| CTCF | ENCSR000ENJ | ENCSR000DTI | yes | 0.9950 | 0.8197 | 0.6316 | 0.8486 |
| CTCF | ENCSR000ENN | ENCSR000DTR | yes | 0.9932 | 0.7788 | 0.5621 | 0.8098 |
| CTCF | ENCSR000EPP | ENCSR000DWQ | yes | 0.9939 | 0.7720 | 0.5517 | 0.7975 |
| CTCF | ENCSR000EOR | ENCSR000DVQ | yes | 0.9939 | 0.8048 | 0.6094 | 0.8319 |
| CTCF | ENCSR000EPR | ENCSR000DPM | no | 0.9913 | 0.7322 | 0.4834 | 0.7600 |
| CTCF* | ENCSR000EOQ | ENCSR000DLW | yes | 0.9962 | 0.7270 | 0.4030 | 0.7868 |
| JUN* | ENCSR000EOQ | ENCSR000EFA | yes | 0.9965 | 0.631 | 0.1644 | 0.6996 |
| MAX* | ENCSR000EOQ | ENCSR000EEZ | yes | 0.9967 | 0.4004 | 0.0255 | 0.3327 |
| MYC* | ENCSR000EOQ | ENCSR000DLU | yes | 0.9977 | 0.1989 | 0.000 | 0.0336 |

**Supplementary Table S6:**
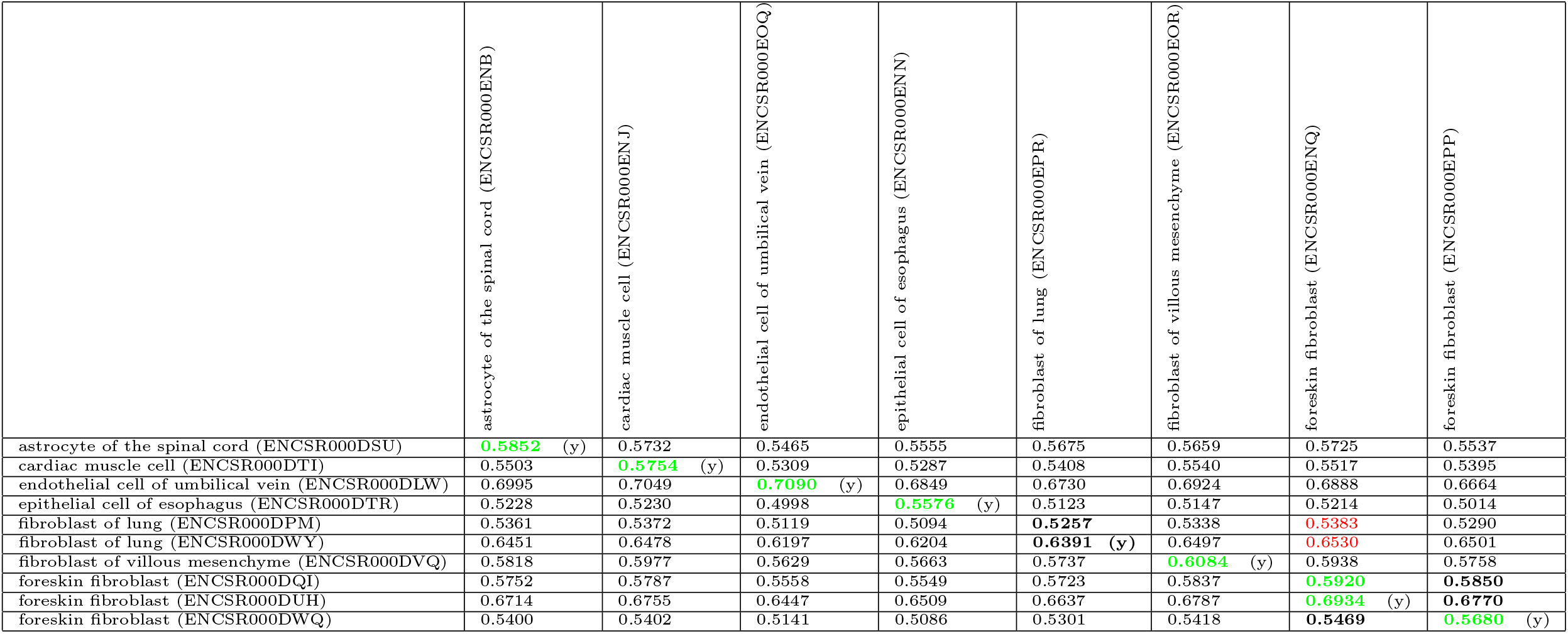
Jaccard coefficients between predicted (columns) and experimentally determined (rows) peak files for CTCF. Entries of matching tissues/cell types are marked in bold. In each row, we mark the largest value in green for matching cell types and in red for differing cell types. We mark matching donor with “(y)”· Jaccard coefficients are computed using the `intersect` and `union` of the GenomicRanges R package. For each peak list, entries are sorted by score and limited to the minimum number of peaks across all peak lists. We find that many of the cell type-specific predictions for CTCF are more similar to the ChIP-seq peaks determined for “endothelial cells of umbilical vein” than to those of their cell type of origin according to the DNase-seq data. One reason might be that only for this experiment (ENCSROOODLW), peaks have not been called using the uniform ENCODE pipeline including SPP (Kharchenko *et al*., 2008), but by another, “unknown” software. However, if we in turn ask for each experimentally determined peak list, which of the predicted peak lists is the most similar one, this picture becomes more encouraging. For 7 of the 8 cell types with matching donor between ChIP-seq and DNase-seq data, the most similar prediction is obtained for the true cell type, while in one case (“fibroblast of lung”), the most similar cell type is “foreskin fibroblast”.

**Supplementary Table S7:**
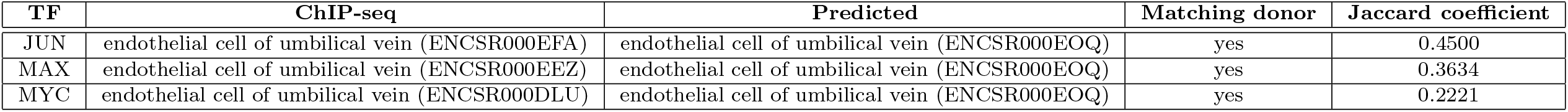
Jaccard coefficient between experimentally determined and predicted peak files. Jaccard coefficients are computed using the `intersect` and `union` of the GenomicRanges R package. For each TF, entries of the experimentally determined and predicted peak lists are sorted by score and limited to the minimum number of peaks in either of the two peak lists.

**Supplementary Figure S6:**
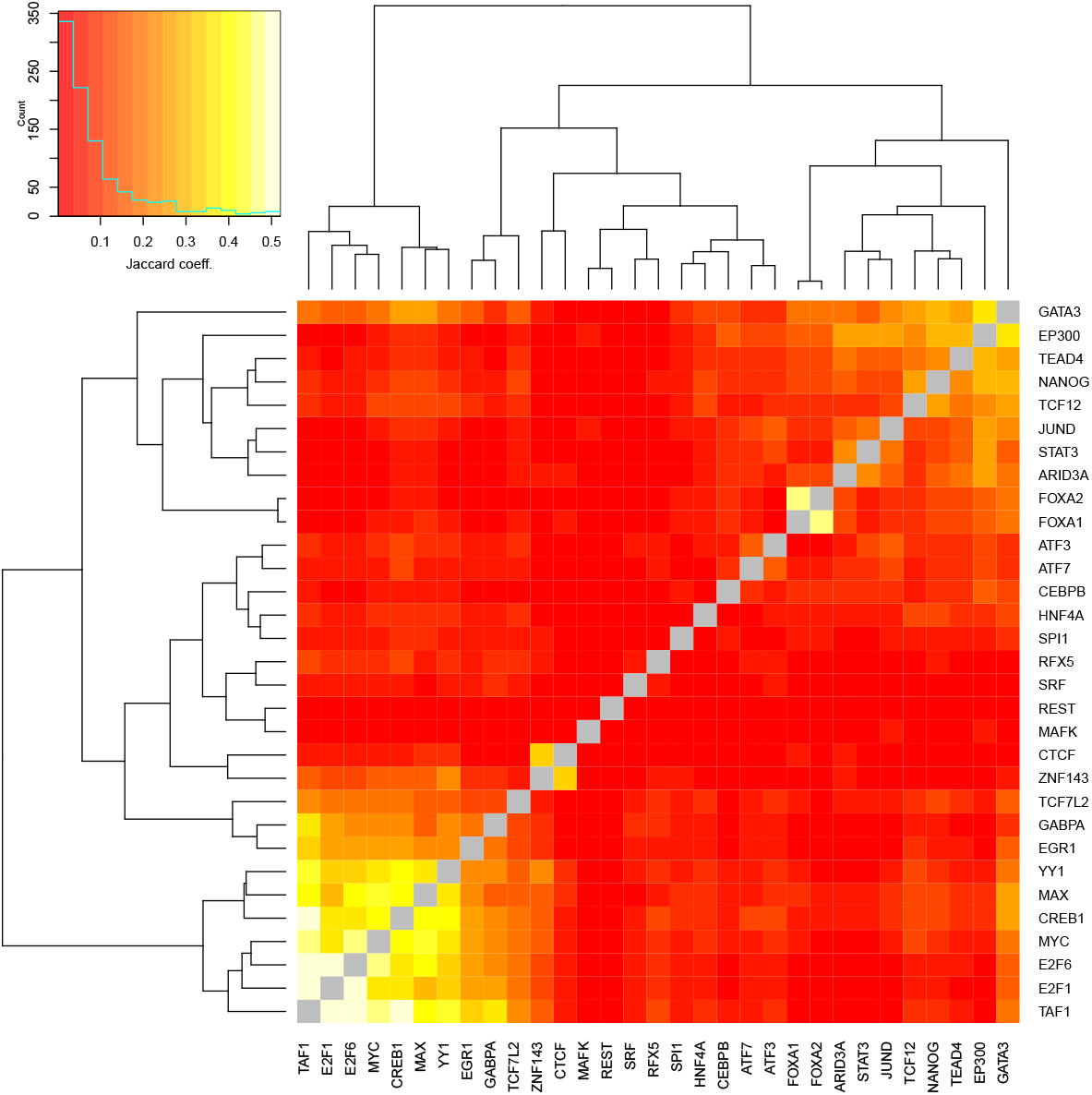
Average Jaccard coefficients computed on the overlap of the peak files of pairs of TFs for matched cell types. In the color scale, the solid cyan line represents the histogram of values observed in the heatmap. Dashed lines indicate the value at the center bin of the color scale. Rows and columns are clustered by the R hclust function using complete linkage.

**Supplementary Table S8:** Benchmark of the simplified open source implementation (Catchitt) of the presented approach compared with the challenge implementation using only DNase-based and motif-based features for the within cell type case (cf. Supplementary Figure S2). For each of the TFs considered, we report AUC-PR achieved by the challenge implementation using only DNase-based and motif-based features (“DNase+motif”), the open source Catchitt implementation. For Catchitt, we additionally consider using only the classifier of the first iteration (in analogy to the comparison in Supplementary Figure S3). We also list the number of motifs utilized in the respective runs for a specific TFs. For the Catchitt runs, we deliberately limited the number of motifs considered to approximate a real-world application of the software. We finally report the ranks among the challenge participants according to the results available at https://www.synapse.org/#!Synapse:syn6131484/wiki/412905.

| TF | cell type | DNase+motif |  |  | Catchitt |  |  |  |
| --- | --- | --- | --- | --- | --- | --- | --- | --- |
|  |  | AUC-PR | Rank | #motifs | AUC-PR | Rank | #motifs | first iteration |
| CTCF | IPSC | 0.810 | 4 | 177 | 0.776 | 6 | 13 | 0.760 |
| NANOG | IPSC | 0.489 | 2 | 127 | 0.404 | 4 | 13 | 0.366 |
| FOXA1 | liver | 0.544 | 2 | 121 | 0.465 | 4 | 12 | 0.435 |
| HNF4A | liver | 0.671 | 4 | 123 | 0.615 | 6 | 10 | 0.597 |
| TAF1 | liver | 0.473 | 4 | 160 | 0.412 | 6 | 12 | 0.400 |

## Supplementary Methods

### Supplementary Text S1 – Tools for predicting in-vivo binding regions

Most approaches (e.g., Pique-Regi *et al*. (2011); Natarajan *et al*. (2012); Piper *et al*. (2013); Gusmao *et al*. (2014); Chen *et al*. (2017)) use binding motifs represented as position weight matrix (PWM) models that have been obtained from databases like TRANSFAC (Matys *et al*., 2006), Jaspar (Mathelier *et al*., 2016), UniProbe (Newburger and Bulyk, 2009) or CisBP (Weirauch *et al*., 2014), or from motif collections like Factorbook (Wang *et al*., 2012), the ENCODE-motif collection (Kheradpour and Kellis, 2014), or Homer (Heinz *et al*., 2010), while some perform de-novo motif discovery based on k-mers (Arvey *et al*., 2012) or as part of convolutional neural networks (Quang and Xie, 2017; Qin and Feng, 2017). Irrespective of the source of the motifs considered, three general schemas are have been established for combining motif predictions with chromatin accessibility data. First, motif matches (i.e., predicted binding sites) may be used as prior information and combined with DNase-seq data to distinguish functional from non-functional binding sites (e.g., Pique-Regi *et al*. (2011); Jankowski *et al*. (2016); Raj *et al*. (2015)), Second, TF footprints may be first identified from DNase-seq data and then annotated with specific TFs based on motif matches afterwards (Gusmao *et al*., 2014). Third, both sources of information are combined in a holistic approach (Quang and Xie, 2017; Qin and Feng, 2017). DNase-seq (and ATAC-seq) data are employed in different ways by existing approaches including i) binning of chromatin accessibility statistics in larger genomic regions around putative binding sites (Luo and Hartemink, 2012), ii) association of chromatin accessibility with specific genes (Schmidt *et al*., 2017), or iii) high-resolution maps of DNase cut sites (Sherwood *et al*., 2014; Raj *et al*., 2015), which may additionally be considered separately for each DNA strand (Piper *et al*., 2013). On the methodological level, approaches either follow a supervised approach based on training examples labeled as “bound” or “unbound”, typically derived from TF ChIP-seq data (e.g., Arvey *et al*. (2012); Luo and Hartemink (2012); Kahara and Lahdesmäki (2015); Liu *et al*. (2017)), or an unsupervised approach clustering regions into “bound” and “unbound” based on their experimental properties (e.g., DNase-seq data or histone modifications (Pique-Regi *et al*., 2011; Sherwood *et al*., 2014; Gusmao *et al*., 2014)), while others base their predictions on statistical tests (Piper *et al*., 2013) or scores related to binding affinity predictions (Schmidt *et al*., 2017). Supervised approaches use a variety of methods like support vector machines (Arvey *et al*., 2012; Kumar and Bucher, 2016), (sparse) logistic regression (Natarajan *et al*., 2012; Luo and Hartemink, 2012; Kahära and Lähdesmaki, 2015; Chen *et al*., 2017), random forests (Liu *et al*., 2017), or neural networks adapted by deep learning (Quang and Xie, 2017; Qin and Feng, 2017). Unsupervised approaches use hierarchical mixture models (Pique-Regi *et al*., 2011), hierarchical multi-scale models (Raj *et al*., 2015), hidden Markov models (Gusmao *et al*., 2014), or other probabilistic models (Sherwood *et al*., 2014). In some approaches, sequence-based features besides motif matches (Kumar and Bucher, 2016; Gusmao *et al*., 2014; Chen *et al*., 2017), sequence conservation (Kumar and Bucher, 2016; Liu *et al*., 2017; Chen *et al*., 2017), or additional experimental data like histone modification (Pique-Regi *et al*., 2011; Arvey *et al*., 2012; Gusmao *et al*., 2014) are included into the model. Finally, a subset of approaches uses the prediction of TF binding regions as an intermediate step for predicting gene regulation (Natarajan *et al*., 2012) or tissue-specific gene expression (Schmidt *et al*., 2017).

### Supplementary Text S2 – Features

The features described in the following are all determined on the level of genome bins. We refer to the bin for which the a-posteriori probability of being peak center should be computed (i.e., the bin containing the peak summit in case of positive examples) as *center bin.* Further, adjacent bins considered are defined relative to that center bin (see also section Prediction schema).

#### S2.1 Sequence-based features

As a first sequence-based feature, we consider the raw DNA sequence according to the *hg19* human genome sequence in the center bin and the directly preceding and the directly following bin. In total, this corresponds to 150 bp of sequence, centered at the center bin.

We further consider the mean G/C-content, and the relative frequency of CG dinucleotides in the raw sequence spanning those three bins centered at the center bin. G/C-content might be an informative property of promoters bound by a certain TF, and an enrichment of CG di-nucleotides might be informative about the presence of CpG islands.

We also compute the Kullback-Leibler divergence between the relative frequencies of all tri-nucleotides in each of these three bins compared with their relative frequencies in the complete genome. As a feature, we then consider the maximum of those three Kullback-Leibler divergence values obtained for the three bins. Here, the reasoning is that a deviation from the genomic distribution of tri-nucleotides might be a sign of the general information content of a sequence, which might help to distinguish coding and non-coding DNA regions as well as identifying regions that encode regulatory information.

Finally, we consider the length of the longest poly-A or poly-T tract, the length of the longest poly-C or poly-G tract, the length of the longest poly-A/T tract, and the length of the longest poly-G/C tract in these three bins.

All of those sequence-based features are neither TF-specific nor cell type-specific, but model parameters learned on their feature values might well be different for different training TFs or cell types.

#### 52.2 Annotation-based features

Based on the Gencode v19 genome annotation of the hg19 genome, we derive a set of annotation-based features. First, we consider the distance of the current center bin to the closest TSS annotation (regardless of its strand orientation), which might be informative about core promoter regions. Second, we collect the binary information if the current center bin overlaps with annotations of i) a CDS, ii) a UTR, iii) an exon, iv) a transcript, or v) a TSS annotation, separately for each of the two possible strand orientations. Like some of the previous features, this helps to identify coding, non-coding but transcribed, core promoter, and intergenic regions. Again, these features are not TF or cell type-specific, but model parameters may be adapted specifically for a TF or cell type.

#### Motif-based features

As it might be expected that binding motifs are pivotal for predicting TF-specific binding regions, we create a large collection of motifs for each of the TFs considered. For each of the TFs, we collect all position weight matrix models from the HOCOMOCO database (Kulakovskiy *et al*., 2016) as well as our in-house database DBcorrDB (Grau *et al*., 2015a), and Slim/LSlim models of the respective TFs from a previous publication (Keilwagen and Grau, 2015). In addition, we learn a large set of motifs from the data provided in the challenge using our motif discovery tools Dimont (Grau *et al*., 2013) using PWM as well as LSlim(3) models (Keilwagen and Grau, 2015). Specifically, we perform motif discovery for

- PWM models from the “conservative” peak files for each training cell type,
- PWM models from the “relaxed” peak files complemented by negative regions selected to be DNase positive (i.e., open chromatin) but ChIP-seq negative according to the ChIP-seq and DNase-seq peak files provided with the challenge data,
- LSlim(3) models from the “conservative” peak files for each training cell type,
- LSlim(3) models from the “relaxed” peak files for each training cell type,
- LSlim(3) models from the “relaxed” peak files complemented by negative regions selected to be DNase positive (i.e., open chromatin) but ChIP-seq negative according to the ChIP-seq and DNase-seq peak files provided with the challenge data.

LSlim(3) may capture intra-motif dependencies between binding site position with a distance of at most three nucleotides.

Motifs discovered using models of different complexity on these different sets of training data (“conservative” and “relaxed” peaks, and “relaxed” peaks complemented by DNase positive regions) should i) capture the breadth of the binding landscape of a TF as represented by the different levels of stringency (“conservative” vs. “relaxed”), and ii) represent potential intra-motif dependencies as well as traditional, “additive” binding affinities. In addition, we learn motifs from the DNase-seq peak files as well, considering

- LSlim(3) models from the “conservative” and “relaxed” DNase-seq peak files,
- LSlim(3) models from the regions in the intersection of all “relaxed” DNase-seq peak files.

Learning motifs from the DNase-seq data alone might have the potential to capture additional binding motifs of TFs that are important for cell type-specific predictions but are not represented in the ChIP-seq data provided with the challenge data.

Regardless of the TF considered, we further include PWM and Slim/LSlim motifs discovered previously (Keilwagen and Grau, 2015; Grau *et al*., 2015a) for CTCF, SP1, JUND, and MAX, as those i) mark boundaries between regulatory regions, ii) frequently interact with other transcriptions factor, or iii) bind to a large fraction of active promoters. Further TFs that might interact with the currently considered TF as determined i) from the literature, specifically from Factorbook (Wang *et al*., 2012), ii) determined from the overlap between the ChIP-seq peaks provided with the challenge data. The latter is accomplished by computing for each TF and cell type i) the TF with the largest overlap (F1 measure computed on the peaks) and ii) the TF with the lowest overlap between the peak files. The former might be indicative of co-binding, while the latter might indicate mutually exclusive binding, both of which might help to predict TF-specific binding regions.

Finally, we consider motifs determined by the epigram pipeline (Whitaker *et al*., 2015), which mark epigenetic modifications. Specifically, we select the top 10 motifs reported for “single mark” analyses for methylation, and H3K4me3 and H3K27ac histone modifications (downloaded from http://wanglab.ucsd.edu/star/epigram/mods/indexhtml).

We use all motif models described above to scan the hg19 genome for potential binding regions. To this end, we apply a sliding window approach across the genome, and aggregate the motif scores obtained according to the genomic bins. For the TF-specific motifs obtained by de-novo motif discovery from ChIP-seq data, we consider as features

- the maximum log-probability of all sliding windows starting in the center bin,
- the logarithm of the sum of binding probabilities in all sliding windows starting in the center bin or its two adjacent bins, and
- the logarithm of the sum of binding probabilities in all sliding windows starting in any of the bins considered.

The first feature should capture the binding affinity at the strongest binding site around the peak summit, while the latter two features represent the general binding affinity of a region with different levels of resolution.

For all of the remaining motifs, we consider the maximum of the bin-wise logarithm of the sum of binding probabilities over all bins considered (see section Binning the genome), as this reduces memory requirements as well as model complexity and this level of detail might be sufficient to capture TF interactions.

#### S2.4 DNase-based features

For the DNase-seq data, the challenge provided tracks with a “fold-enrichment coverage” track, peak files, and the original BAM files from mapping the DNase-seq reads, of which we consider only the former two. From the fold-enrichment coverage track, we compute the following statistics:

- the minimum value across the center bin and its two adjacent bins,
- the minimum of the maximum value within each bin considered,
- the minimum of the 25% percentile within each bin considered, and
- the median values of all the bins considered.

After extracting those feature values for all genomic bins, we quantile normalize each of the features independently across the challenge cell types. Before normalization, we randomize the order of values to avoid systematic effects due to genomic order, which might especially occur for the large number of very low values. For the additional, primary cell types, we do not perform an independent quantile normalization but instead map the DNase-seq features (according to their numerical order) to the corresponding, quantile normalized values of the challenge cell types.

In addition to these short-range DNase features, we also determine a set of long-range features, which are computed from i) 10 bins ii) 20 bins, and iii) 40 bins preceding and succeeding the current center bin. These features are

- the minimum value across all bins,
- the maximum value across all bins,
- the minimum value across the bins preceding the center bin,
- the minimum value across the bins succeeding the center bin,
- the maximum value across the bins preceding the center bin, and
- the maximum value across the bins succeeding the center bin.

Together, these features capture chromatin accessibility on a short and long range level with reasonable resolution, which should be highly informative with regard to the general TF-binding potential. Model parameters should then be able to adapt for TF-specific preferences of chromatin accessibility.

For the current center bin, we additionally determine features of stability across the different cell types, namely

- the ratio of the minimum value in the current cell type divided by the average of the minimum values across all cell types,
- the ratio of the maximum value in the current cell type divided by the average of the maximum values across all cell types,
- the coefficient of variation (standard deviation divided by mean) of the minimum values across all cell types, and
- the coefficient of variation of the maximum values across all cell types,

where the latter two features are identical for all cell types by design.

We also determine several features that represent the monotonicity/stability of these DNase-seq signals. Specifically, these features are

- the number of steps (increasing or decreasing) in the track profile in a 450 bp interval centered at the center bin,
- the longest strictly monotonically increasing stretch in the four bins preceding the center bin,
- the longest strictly monotonically decreasing stretch in the four bins preceding the center bin,
- the longest strictly monotonically increasing stretch in the four bins succeeding the center bin, and
- the longest strictly monotonically decreasing stretch in the four bins succeeding the center bin.

The first of these features has been inspired by the “orange” feature coined by team autosome.ru in the challenge.

Finally, we define further features based on the “conservative” and “relaxed” DNase-seq peak files as provided with the challenge data. These are

1. the distance of the center bin to the summit of the closest conservative peak,
2. the distance of the center bin to the summit of the closest relaxed peak,
3. the peak statistic of a conservative peak overlapping the center bin (or zero if no such overlapping peak exists) multiplied by the length of the overlap,
4. the peak statistic of a relaxed peak overlapping the center bin (or zero if no such overlapping peak exists) multiplied by the length of the overlap,
5. the maximum of the q-values of an overlapping conservative peak (or zero if no such overlapping peak exists) multiplied by the length of the overlap across the five central bins,
6. the maximum of the q-values of an overlapping relaxed peak (or zero if no such overlapping peak exists) multiplied by the length of the overlap across the five central bins.

#### S2.5 RNA-seq-based features

The RNA-seq data provided with the challenge data included the TPM values of genes according to the gencode v19 genome annotation. TPM values are also quantile normalized across the cell types. As features, we consider

- the maximum TPM value (averaged over the two bio-replicates per cell type) of genes in at most 2.5 kb distance
- the coefficient of variation of the bio-replicated of the corresponding gene,
- the relative difference (difference of values in bio-replicated divided by their mean value) of the corresponding gene.

In analogy to the DNase-based features, we computed from the first feature as measures of stability across the different cell types

- the ratio of the maximum TPM value in the current cell type divided by the average of the maximum values across all cell types, and
- the coefficient of variation of the maximum TPM values across all cell types.

### Supplementary Text S3 - Model & learning principle

For numerical features *x*, we use independent Gaussian densities parameterized as

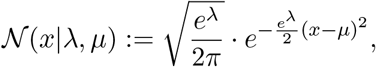

which allows for unconstrained numerical optimization of both, λ and *μ*.

For features *y* with *K* possible discrete values *v*_1_,…, *v_K_*, we use (unnormalized) multinomial distributions with parameters *β* = (*β*_1_,…,*β_K_*) defined as

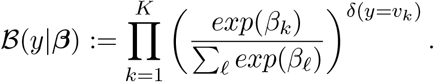

The multinomial coefficient is neglected in this case, since it only depends on the input data but not on the model parameters. In case of binary features, i.e., K=2, this corresponds to an (unnormalized) binomial distribution.

For modeling the raw sequence *s* = *s*_1_ *s*_2_…*s_L_*, *s_ℓ_* ∈ Σ = {*A, C, G,T*}, we use a homogeneous Markov model of order 3 parameterized as

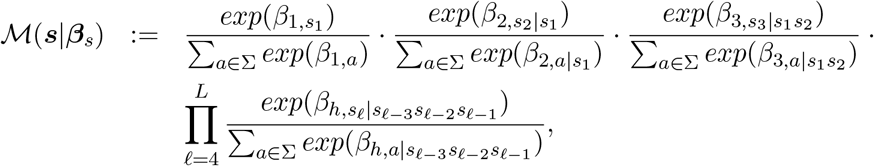

where *β*_*h,a*|*b*_, *a* ∈ Σ, *b* ∈ Σ^3^ are the homogeneous parameters and *β_s_* = (*β*_1,*A*_,…, *β*_1,*T*_, *β*_2,*A*|*A*_,…, *β*_2,*T*|*T*_, *β*_3,*A*|*AA*_,…,*β*_3,*T*|*TT*_,…, *β*_*h*,*A*|*AAA*_,…, *β*_*h,A*|*AAA*_,…, *β*_*h,T*|*TTT*_) denotes the vector of all model parameters.

Let *x* = (*x*_1_,…, *x_N_*) denote the vector of all numerical features, *y* = (*y*_1_,…, *y_M_*) denote the vector of all discrete features, and *s* denote the raw sequence of one region represented by its feature values *z* = (*x, y, s*). Let *θ* = (λ_1_,…, λ_*N*_, *μ*_1_,…, *μ_N_*, *β*_1_,…, *β_M_*, *β_s_*) denote the set of all model parameters. We compute the likelihood of *z* as an independent product of the terms for the individual features, i.e.,

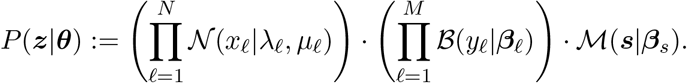

For modeling the distribution in the positive (foreground) and negative (background) class, we use likelihoods *P*(*z*|*θ_fg_*) and *P*(*z*|*θ_bg_*) with independent sets of parameters *θ_fg_* and *θ_bg_*, respectively. In addition, we define the a-priori class probabilities as 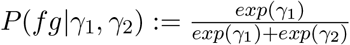 and 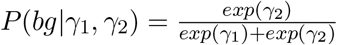.

Based on these definitions, we may compute the a-posteriori class probability of the positive class as

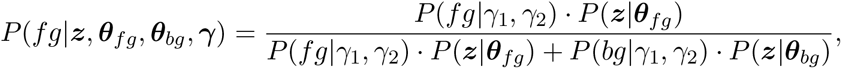

and the a-posteriori class probability of the negative class in complete analogy.

Using the discriminative maximum conditional likelihood principle (Roos *et al*., 2005), the parameters are optimized such that the a-posteriori probabilities of the correct class labels given data and parameters are maximized. Here, we use a variant (Grau, 2010) of the maximum conditional likelihood principle that incorporates weights. Let *F* = (*z*_1_,…, *z*_*I*_) denote the set of positive examples and let *B* = (*z*_*I*+1_,…, *z_J_*) denote the set of negative examples, where *z_i_* is assigned weight *w_i_*. The parameters are then optimized with regard to

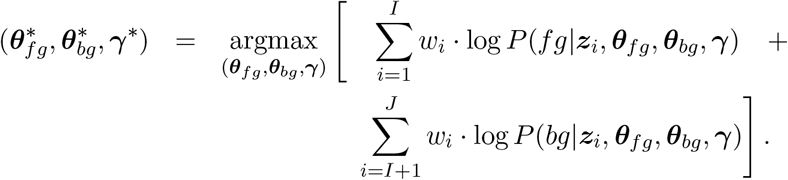

### Supplementary Text S4 - Sampling of DNase-matched negative regions

We sample negative regions with chromatin accessibility values matched to the positive regions (following an idea related to importance sampling) as explained in the following. We consider the center bins of all positive regions, collect the corresponding DNase-seq median feature values (see Supplementary Text S2) of those bins, and determine a histogram of the collected values. The histogram is composed of 20 equally sizes bins between the observed maximum and minimum values of the DNase-seq median values. This histograms represents an approximation of the distribution of DNase-seq median values in the positive regions. As we expect DNase-seq values to be highly informative about TF binding, we aim at sampling a representative set of negative regions that exhibit similar DNase-seq values but might be distinguished from positive regions by other features.

To this end, we assign each of the negative regions to the same histogram bins based on their respective DNase-seq median values at their center bins. This also yields an analogous histogram of the DNase-seq median values for the negative regions, which will usually be different from the histogram for the positive regions.

Within each histogram bin, we then draw a subset of the negative regions assigned to that bin by i) drawing a subset of these regions four times as large as the corresponding positive set, and ii) weighting the drawn negative regions such that the sum of weights matches the relative abundance of that histogram bin in the histogram on all negative region.

Conceptually, this procedure yields an over-sampling of negative regions with large DNase-seq median features, which is adjusted for by down-weighting such examples to the corresponding frequency on the chromosome level. This is especially important as these will be regions that are hard to classify using DNase-seq based features but are only lowly represented by the uniform sampling schema.

